# Altered O-glycosylation Level of SARS-CoV-2 Spike Protein by Host O-glycosyltransferase Strengthens Its Trimeric Structure

**DOI:** 10.1101/2021.04.06.438614

**Authors:** Zhijue Xu, Xin Ku, Jiaqi Tian, Han Zhang, Jingli Hou, Can Zhang, Jingjing Shi, Yang Li, Hiroyuki Kaji, Sheng-Ce Tao, Atsushi Kuno, Wei Yan, Lin-Tai Da, Yan Zhang

## Abstract

The trimeric spike protein (S) mediates host-cell entry and membrane fusion of SARS-CoV-2. S protein is highly glycosylated, whereas its O-glycosylation is still poorly understood. Herein, we site-specifically examine the O-glycosylation of S protein through a mass spectrometric approach with HCD-triggered-ETD model. We identify 15 high-confidence O-glycosites and at least 10 distinct O-glycan structures on S protein. Peptide microarray assays prove that human ppGalNAc-T6 actively participates in O-glycosylation of S protein. Importantly, the upregulation of ppGalNAc-T6 expression can profoundly enhance the O-glycosylation level by generating new O-glycosites and increasing both O-glycan heterogeneity and intensities. Further molecular dynamics simulations reveal that the O-glycosylation on the protomer-interface regions, which are mainly modified by ppGalNAc-T6, can potentially stabilize the trimeric S protein structure. Our work provides deep molecular insights of how viral infection harnesses the host O-glycosyltransferases to dynamically regulate the O-glycosylation level of the viral envelope protein responsible for membrane fusion.

## INTRODUCTION

The novel coronavirus disease 2019 (COVID-19) caused by severe acute respiratory syndrome coronavirus 2 (SARS-CoV-2) has presented a pandemic threat to global public health(Chan et al., 2020b; Chen et al., 2020). SARS-CoV-2 belongs to the family of *Coronaviridae*, same as SARS-CoV and Middle east respiratory syndrome (MERS), etc(Chan et al., 2020a). Similar to SARS-CoV and MERS, the viral surface spike protein (S protein), highly glycosylated, is responsible for mediating the membrane fusion between virus and host cells(Lan et al., 2020). S protein is a homotrimeric type I membrane protein and contains two major subunits, the receptor-binding subunit S1 and the membrane fusion subunit S2(Cai et al., 2020; Wang et al., 2018). It has been proposed that two trimeric S proteins can simultaneously bind to one angiotensinconverting enzyme 2 (ACE2) homodimer from host cells(Yan et al., 2020), and then mediate the membrane fusion process(Hoffmann et al., 2020). Recently, S protein was also found to recognize several immune cell receptors other than ACE2, including MR/CD206 and DC-SIGN/CD209, etc(Gao et al., 2020). Notably, these receptors belong to C-type lectin or mannose receptors, suggesting that S protein could mediate the viral entry in a glycan-dependent manner via its surface glycans(Gao et al., 2020).

S protein is coated with 22 N-glycan sites and numerous O-glycan sites on each monomer(Casalino et al., 2020; Zhao et al., 2020a). The former has been well characterized and was determined to be structurally heterogenous and certain N-glycans are directly involved in regulating the interactions between S protein and ACE2(Casalino et al., 2020; Zhao et al., 2020a). for example, deletions of the N-glycosites on N331 and N343 could drastically reduce the viral infectivity(Li et al., 2020b). The O-glycosylation of S protein was also previously characterized using mass spectrometry (MS) analysis(Sanda et al., 2021; Shajahan et al., 2020; Watanabe et al., 2020; Zhang et al., 2020; Zhao et al., 2020a). Nevertheless, most of the former studies used higher-energy collisional dissociation (HCD) mode to fragment the O-glycopeptides, which can cause significant sugar losses and is difficult to pinpoint the accurate O-glycosites(Vakhrushev et al., 2013). The electron-transfer dissociation (ETD) mode, however, formats the fragment ions containing both glycans and peptides, allowing determining the O-glycosites unambiguously(Lermyte et al., 2018). To date, the ETD fragmentation method has been widely used in dissecting the location and structures of O-glycans for many biomolecules(Hao et al., 2019; Riley et al., 2020; Steentoft et al., 2011).

O-linked glycosylation, also named as the mucin-type O-glycosylation, is one of the most abundant glycosylation in human(Kudelka et al., 2015; Steentoft et al., 2013) and plays critical roles in modulating various functions of O-glycoproteins, including proprotein processing, subcellular location and ligand-receptor interactions et al(Goth et al., 2018; Liu et al., 2017; Wang et al., 2018). The processing of O-glycosylation mostly takes place in Golgi apparatus, and is initially controlled by one glycosyltransferase family of polypeptide N-acetylgalactosaminyltransferases (ppGalNAc-Ts)(Bennett et al., 2012; Li et al., 2012). Currently, up to 20 ppGalNAc-Ts have been discovered in human, each isoenzyme catalyzes distinct but partly overlapping protein substrates(De Las Rivas et al., 2020; Gerken et al., 2011; Schjoldager et al., 2015; Xu et al., 2017). It was reported that the expressions of ppGalNAc-T isoenzymes were changed dramatically in both COVID-19 patients samples and the SARS-CoV-2 infected A549 cells suggesting that the host O-glycosyltransferases were regulated by SARS-CoV-2 infection(Liao et al., 2020; Stukalov et al., 2020). It is noteworthy that the O-glycan modifications also participate in regulating the stability and biological functions of certain viral proteins(Stone et al., 2016). However, the dynamic regulations of O-glycosylation level induced by SARS-CoV-2 infection and the biological effects of the O-glycosylation on SARS-CoV-2 remain unknown.

In this study, we employed an HCD triggered ETD MS approach to characterize the O-glycan sites/structures for the complete S protein, thereby a total of 15 O-glycosites are identified and for each site, the glycan structures are determined or inferred. Importantly, increase ppGaNAc-T6 expression can lead to pronounced altered O-glycan patterns on S protein, which in turn favors the formation of its trimeric structure as revealed by molecular dynamics (MD) simulations. Our work provides deep structural insights into the functional role of certain host O-glycosyltransferase in dynamically regulating the O-glycosylation level of the viral glycoproteins, thereby facilitate the membrane fusion.

## RESULTS

### SARS-CoV-2 S protein is both N- and O-glycosylated in human HEK-293T cell

We expressed a recombinant ectodomain of S protein (S-ECD) in the human embryonic kidney 293T (HEK-293T) cells (Figure 1A). To prevent the furin cleavage of the S protein at the ^682^RRAR^685^ site, we introduced a “GAAG” substitution at the above cleavage site and additionally designed two proline mutations at Lys986 and Val987 to increase the stability of the trimeric S protein, as suggested by former work(Pallesen et al., 2017). The S protein from culture medium and cell lysates was purified via immunoprecipitation with anti-FLAG antibody, and the protein purity was analyzed by silver staining (Figure 1B). The S protein purified from culture medium displays smeared bands, however, the protein derived from cell lysate is shown by one sharp band, suggesting that the secreted S protein was highly glycosylated. To examine the nature of the glycans, we treated the S protein with either N-glycosidase PNGase F or neuraminidase. As a result, both treatments can lead to profound decrease of the protein molecule-weight (Figure 1C), confirming that the S protein is coated with N-glycans. Notably, further treatment by O-glycosidase, together with PNGase F and neuraminidase, can give rise to additional molecular-weight drop of S protein comparing to that treated by PNGase F or neuraminidase alone (Figure 1C). The above results indicate that the S protein expressed in HEK-293T cells is both N- and O-glycosylated.

**Figure 1.**
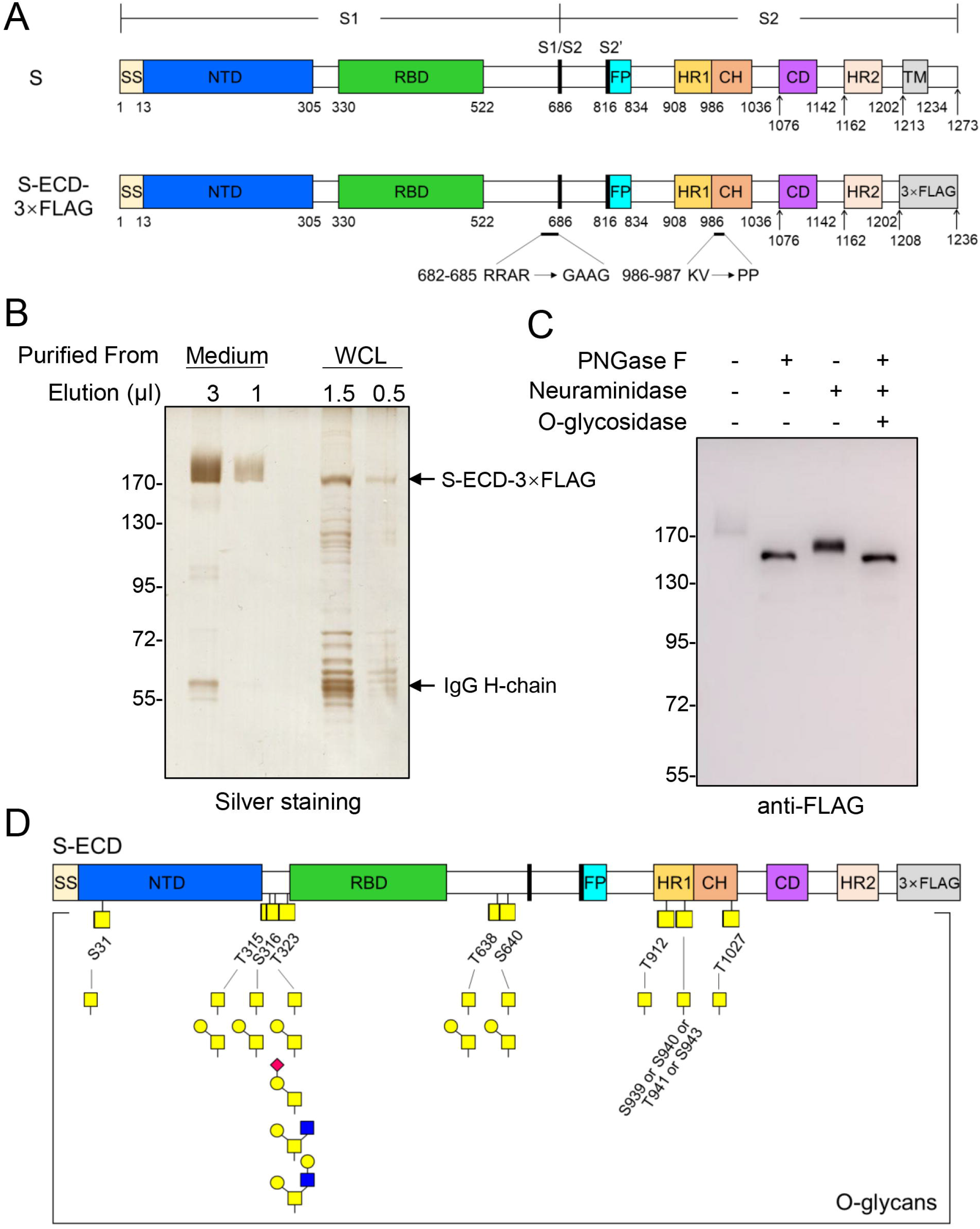
Expression and purification of SARS-CoV-2 S protein. (A) Schematic diagram of SARS-CoV-2 S protein and recombinant S-ECD-3×FLAG protein. The protein domains are illustrated: SS, signal sequence, NTD, N-terminal domain, RBD, receptor-binding domain, FP, fusion peptide, HR1, heptad repeat 1, CH, central helix, CD, connector domain, HR2, heptad repeat 2, TM, transmembrane domain(Watanabe et al., 2020). In recombinant S-ECD-3×FLAG protein, “GAAG” was introduced to replace the cleavage site “RRAR” and the K986-V987 motif were substituted with “PP”. (B) Silver staining of purified S-ECD-3×FLAG protein. The S-ECD-3×FLAG protein was expressed in human embryonic kidney 293T cells, and purified from culture medium or whole cell lysate (WCL) by anti-FLAG beads. (C) Glycosidase treatment of purified S-ECD-3×FLAG protein. The purified S-ECD-3×FLAG proteins (50 ng) were treated with PNGase F, neuraminidase and/or O-glycosidase, respectively. After treatment, the proteins were separated by SDS-PAGE and immunoblotted with anti-FLAG antibody. (D) Identified O-glycosites and corresponding O-glycan structures on S-ECD-3×FLAG protein. See also Figure S1 and Table S1.

### Site-specific analysis of O-glycosylation on SARS-CoV-2 S protein

To pinpoint the O-glycosites and the corresponding glycan structures, we conducted the MS analysis using a sequential fragmentation strategy (HCD triggered ETD). We firstly optimized the MS parameters using a commercialized recombinant S1 subunit as a standard sample. At least 7 high-confidence O-glycosites and 10 O-glycan structures were identified from the S1 subunit (Figure S1 and S2). Using the optimized MS parameters, we then examined the O-glycosites and structures of the S protein purified from HEK-293T cells. As a summary result as shown in Figure 1D, we identified a total of 9 O-glycosites, including 5 Thr-site (T315, T323, T638, T912, and T1027), 3 Ser-site (S31, S316, and S640), and one ambiguous site within the S939-S943 motif. The Tn and T structures are the dominant O-glycan types on S protein, while sialylated T structure (ST), core-2 type GalNAcGal(GlcNAc) glycan (C2), or core-2 type GalNAcGal(GlcNAcGal) glycan (C2-2) were also detected, mostly at T323.

### An on-chip ppGalNAc-Ts assay on S protein peptide microarray

Interesting, in the case of glycosylation processing, proteomics studies have shown that the expression of host glycosyltransferases could be modulated after the SARS-CoV-2 infection(Stukalov et al., 2020). Consistently, in a public available data of single-cell RNA sequencing on bronchoalveolar lavage fluid cells of patients with COVID-19(Liao et al., 2020), the expressions of ppGalNAc-Ts were changed dramatically in patients that further indicated that the host O-glycosyltransferases were regulated by SARS-CoV-2. To further evaluate the functional roles of different ppGalNAc-Ts in S protein O-glycosylation, we examined the glycosyltransferase activities against short peptides derived from S protein using the peptide microarray that contains 211 peptides covering the full-length of S protein (each with 12-aa in length)(Li et al., 2020b). The attached Tn structure was detected using the *Vicia villosa* agglutinin (VVA) (Figure 2A). Four ppGalNAc-T isoenzymes of T1, T2, T3, and T6 were chose for the analysis because they were the major players in human O-glycosylation (Figure 2B)(Kong et al., 2015). The peptides with signal-to-noise ratio larger than 5.0 were considered as positive substrates for each ppGalNAc-T. As a result, twenty-two peptides were enriched from four ppGalNAc-Ts, and the most 15 peptides of them were glycosylated by T6 (Figure 2C). The results clearly show that T6 exhibits high enzymatic activities against a broader set of peptides comparing to other three isoenzymes. Moreover, T1 and T2 prefer to catalyze the flexible regions connecting functional domains, whereas T6 can also target to the structured units, i.e., N-terminal domain (NTD) and connector domain (CD). This microarray assay suggests that T6 is the major ppGalNAc-T involved in the O-glycosylation of S protein.

**Figure 2.**
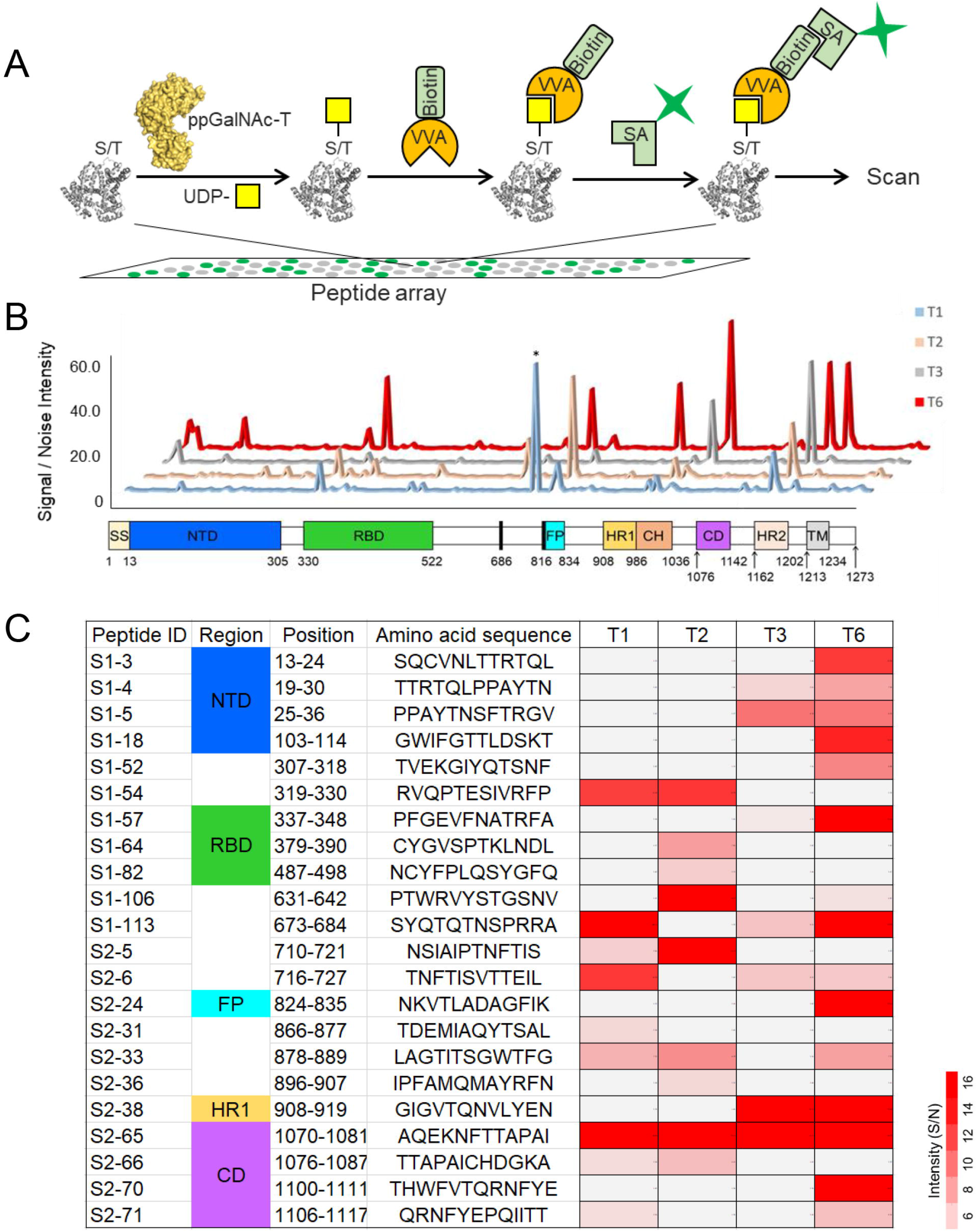
The on-chip ppGalNAc-Ts assay on S protein peptide microarray. (A) Scheme of on-chip ppGalNAc-Ts assay. The peptide microarray contains short peptides covering the full-length of S proteins. The ppGalNAc-T transfers GalNAc from UDP-GalNAc to the targeted Ser/Thr residues, and the O-GalNAc glycan were then immobilized with biotin labeled VVA lectin and Cy5 labeled streptavidin. The signal is readily analyzed by a microarray scanner. (B) An overview of S protein peptide glycosylation modified by T1/T2/T3/T6, and (C) the heatmap of the glycopeptides with a signal to noise (S/N) intensity over 5.0 were shown. The intensity marked with an asterisk is ~180.

### Overexpression of ppGalNAc-T6 dramatically alters the O-glycan patterns of S protein

To further investigate the biological implications of ppGalNAc-T6 on S protein, we expressed and purified the S protein from the T6-coexpressed HEK-293T cells, and then, determined the resulting O-glycan features. Interestingly, increease T6 expression can profoundly alter the O-glycosylation pattern on S protein compared to that obtained from the normal HEK-293T cell. In specific, 5 additional O-glycosites were observed upon T6 upregulation, including three Thr-sites (T29, T307 and T768), one Ser-site (S459) and one uncertained site at T572 or T573. In addition, more diversified O-glycan structures were observed at certain sites (Figure 3A). That is, additional T structure at S31 and T912; ST structure at T315, S316, and T638; C2 structure at T638 were identified.

**Figure 3.**
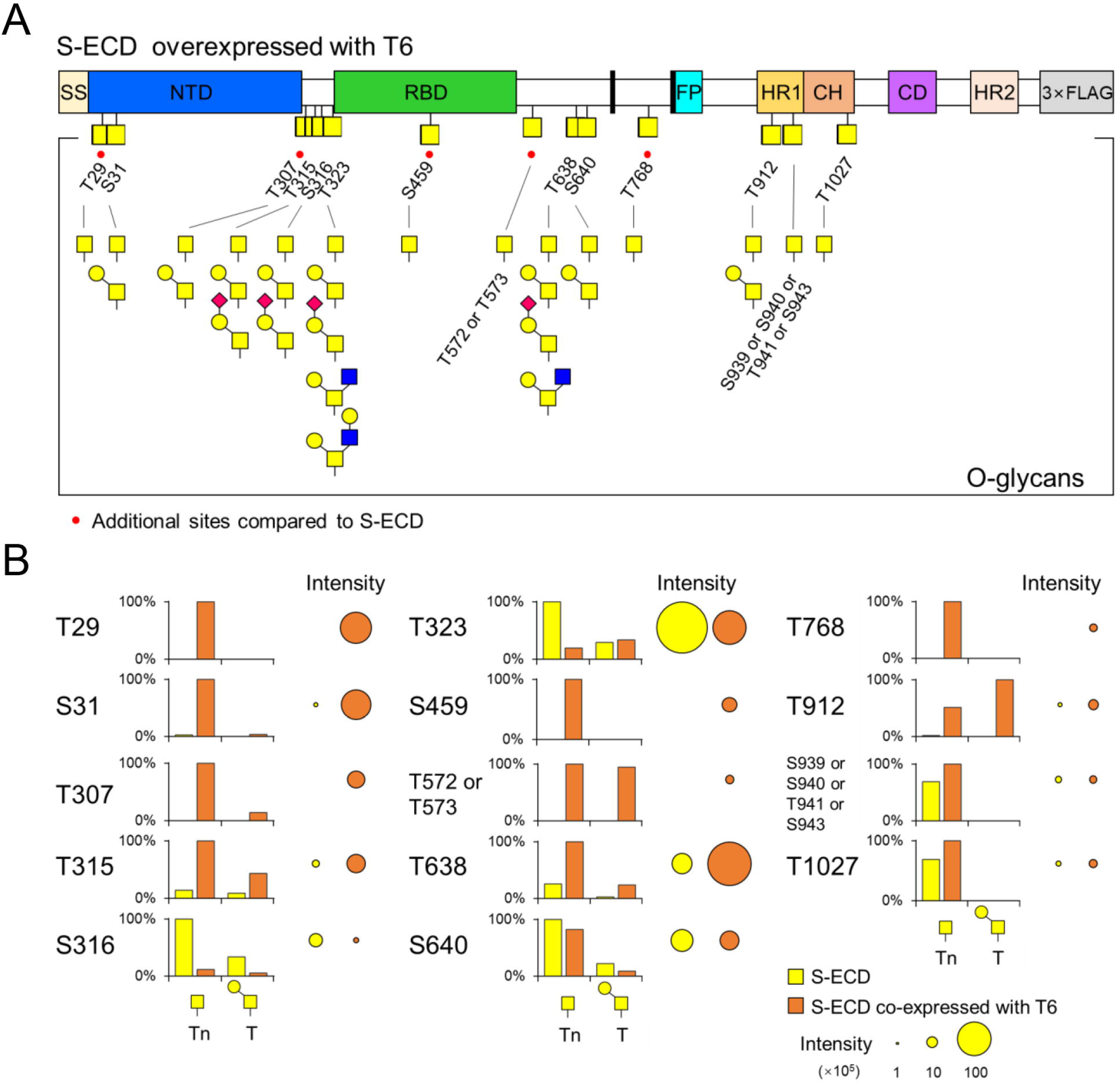
Site-specific O-glycosylation of S protein with overexpressed ppGalNAc-T6. (A) Identified O-glycosites and O-glycans on S-ECD-3×FLAG from the T6-overexpressed HEK-293T cells. The newly discovered O-glycosites upon T6 overexpression are highlighted with red dots. The representative MS/MS spectra were shown in Supplementary Figure 6. (B) Quantitative comparisons of the O-glycosylation level for each identified O-glycosites on S protein. The O-glycan intensities at each site were normalized to the max intensity of the O-glycan at that site. Bubbles indicate the total O-glycan intensity at each site. The relative O-glycan intensities of S-ECD with and without overexpressed T6 were normalized according to the intensities of base peak chromatograms of each sample. See also Table S1.

More intriguingly, we find that the O-glycosylation levels at S31, T315, T638, and T912 are profoundly increased upon T6 coxpression, whereas that at S316 and T323 are significantly decreased. Meanwhile, the O-glycosylation at S640, T1027, and one uncertained site within the S939-S943 motif remains nearly unchanged (Figure 3B). These findings suggest that the enhanced T6 expression can significantly alter the O-glycosylation patterns on S protein.

### The enhanced O-glycosylation on S protein exerted by T6 stabilizes its trimeric conformation

As described above, we identify a total of 15 O-glycosites on one S protein protomer, with 9 glycosites located on S protein surface and the others lying in the interfaces between protomers (Figure 4A and 4B). To examine how the interface O-glycosites might affect the structure of the S protein trimer, we further performed molecular dynamics (MD) simulations for each of the six interface-glycosites with and without Tn modification (Figure 4C). Intriguingly, we find that the O-glycosylation on T1027, T912, T768, T572 and S316 can profoundly promote the formation of the S protein trimer by strengthening the inter-protomer interactions. In specific, for the T1027-site, the Tn-group from three protomers can form stable non-polar contacts via their −CH3 groups, and can also establish one hydrogen bond (HB) with the R1039 sidechain from the neighboring protomer (Figure 4D and S3C). The above interactions stabilize the S-trimer conformation, reflected from the shortened distance between the T1027 Cα atoms from each pair of adjacent protomers (Figure 4D). Likewise, the GalNAc-group on the T912-site can directly contact with F1121 from the adjacent protomer via stacking interactions, which also strengthens the S-trimer conformation (Figure 4E).

**Figure 4.**
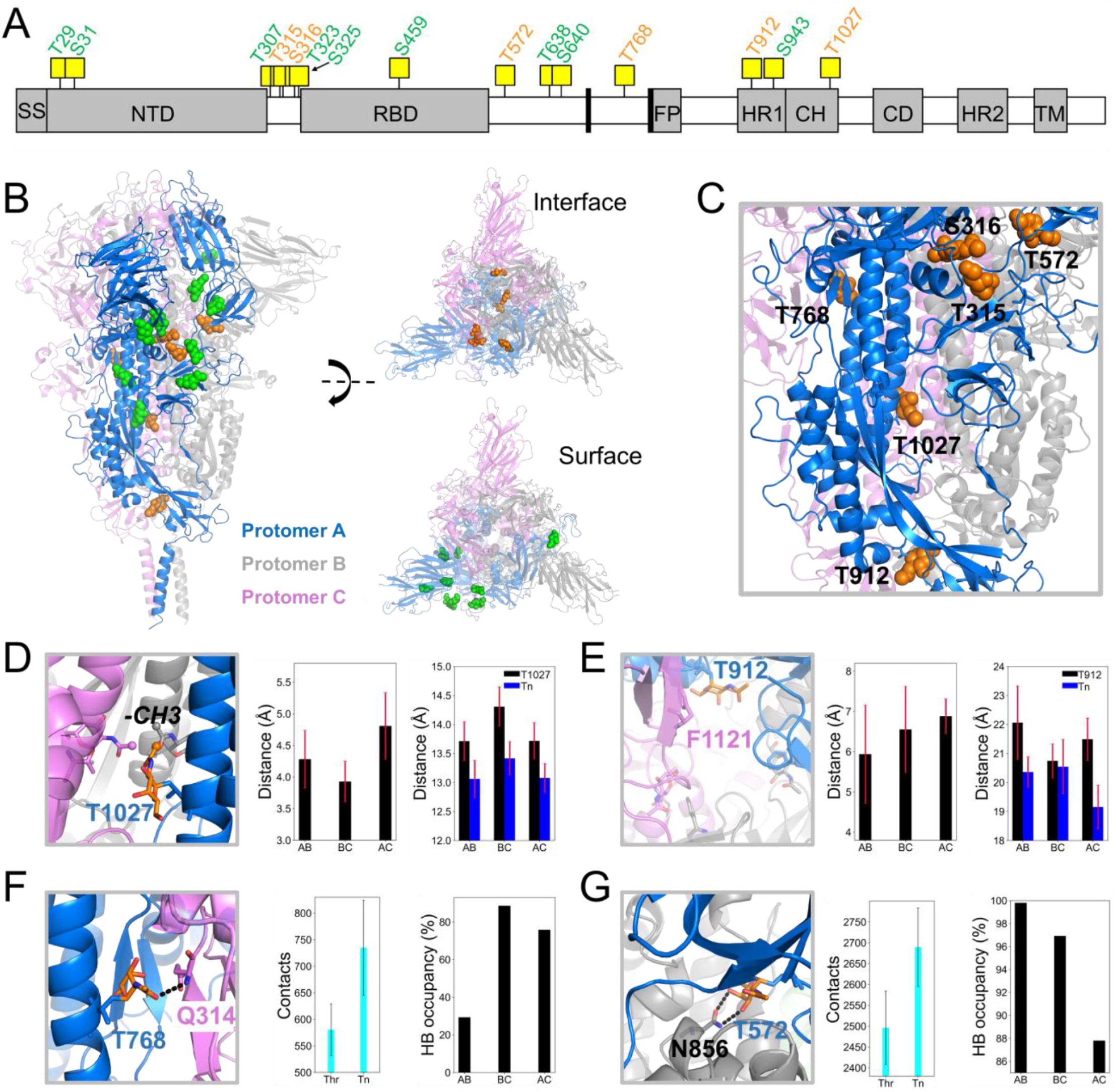
3D modeling and molecular dynamics simulations of O-glycosylation on S protein. (A) A schematic illustration of the S-protein, with 15 identified O-glycosites labeled on top. In particular, the O-glycosites locating on the S protein surface and protomer interfaces are colored in green and orange, respectively. (B) Structural representation of S protein trimer in two different views. Protomer A, B, and C are shown in blue, gray, and violet, respectively. The modeled Tn-structure on protomer A are highlighted in spheres, colored differently according to the sites. (C) A zoomed-in view of 6 O-glycosites locating in the protomer interfaces. (D) The structural highlight of the T1027-Tn from three protomers, with the CH3-group of Tn shown in spheres. Inter-Tn:CH3 distance (middle panel) & T1027 Cα atom distance (right panel) between each two S-protomers. (E) The structural highlight of the T912-Tn from three protomers. F1121 is shown in colored sticks. The distances between the center of mass (COM) of GalNAc and COM of F1121 sidechain from the neighbor protomer (middle panel). The T912 Cα atom distance (right panel) between each two S-protomers. (F) The structural highlight of the T768-Tn in protomer A; Q314 from the protomer C is shown in violet sticks. Comparisons of the inter-protomer contact numbers with and without Tn modifications (Middle panel), the contact number is calculated by sum of each two protomer contacts surrounding the Tn-structure with a distance cutoff of 6 Å; HB occupancy between Tn and Q314 from the neighboring protomer (Right panel). (G) The structural highlight of the T572-Tn in protomer A; N856 from the protomer B is shown in gray sticks. Comparisons of the inter-protomer contact numbers with and without Tn modifications (Middle panel). HB occupancy between T572-Tn and N856 from the neighboring protomer (Right panel). Each HB is highlighted in black dashed line. See also Figure S3.

In addition, for the T768, T572, and S316 sites, the presence of Tn structure can significantly increase the contact number between two adjacent protomers compared to the non-glycosylated conformations (Figure 4F, 4G and S3A). Moreover, the T768-Tn and T572-Tn can also interact with Q314 and N856 from different protomer, respectively, via HB interactions (Figure 4F and 4G). The GalNAc-group on the S316-Tn, on the other hand, can form a water-mediated interaction with T761 from neighboring protomer (Figure S3A). In contrast, the O-glycosylation of S315 imposes no apparent structural perturbations on S protein trimer (Figure S3B). Altogether, most of the interface O-glycosites are found to play critical roles in stabilizing the S protein trimer structure by forming hydrophobic and/or HB interactions with other protomers. Notably, the O-glycosylation levels of three interface O-glycosites, T912, T768, and T572, are greatly enhanced by T6 upregulation, highlighting again the significant role of T6 in modulating the S protein trimeric structure.

### Bioinformatic analyses of the O-glycosites on the S proteins from SARS-CoV and SARS-CoV-2

We next looked into the conservation of the identified 15 O-glycosites among 492,167 SARS-CoV-2 strains collected from the 2019 novel coronavirus resource database of the National Genomics Data Center (NGDC) updated to 2021/02/08(Gong et al., 2020; Song et al., 2020; Zhao et al., 2020b). Our analyses show that the top 5 high mutation-rate sites are D614, A222, I68, 681P, and 501N, among which the D614-site displays a mutation rate of ~60%. In sharp contrast, all the 15 O-glycosites exhibit a mutation rate less than 0.1%, except S939 (~0.16%) (Figure 5A & Table 1), suggesting that the O-glycosites of S protein in SARS-CoV-2 are relatively conserved among the SARS-CoV-2 strains. Moreover, we also examined the O-glycosite differences between the S proteins of SARS-CoV-2 and SARS-CoV (Figure 5B and S4, Table 1). Among the 15 O-glycosites of SARS-CoV-2 S protein, six are neither Ser nor Thr in SARS-CoV S protein. Notably, except T572, all other five O-glycosites are located on the protein surface. Therefore, it can be expected that the interface O-glycosites likely play a common role in stabilizing the S trimer structure among SARS-CoV and SARS-CoV-2.

**Figure 5.**
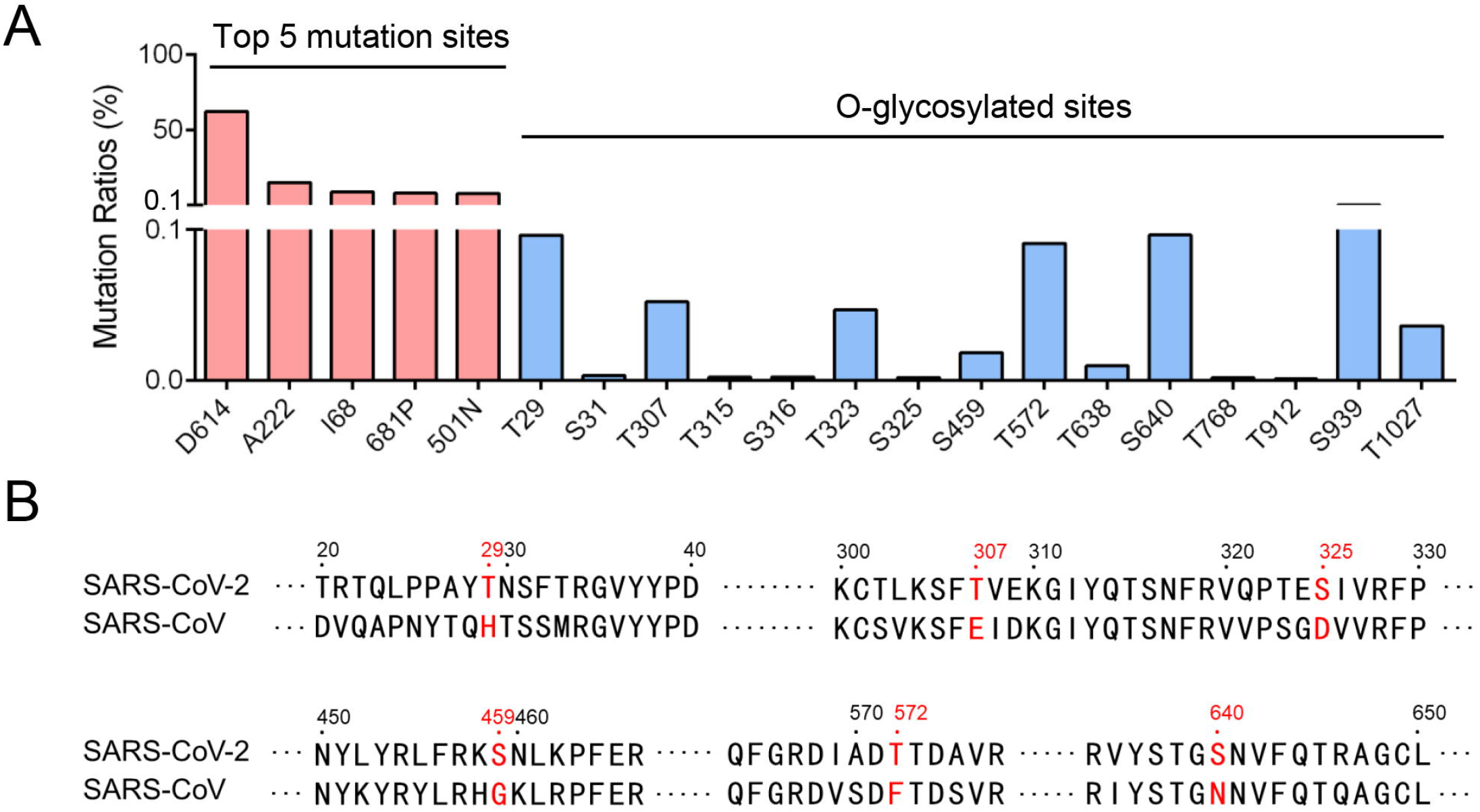
Mutations of the S protein O-glycosites among SARS-CoV-2 strains and comparisons to SARS-CoV. (A) Mutation ratios of the O-glycosites in public available SARS-CoV-2 mutated strains. Top 5 mutation sites of S protein were also listed out and colored in red. The data were referred to the 2019 novel coronavirus resource database of the National Genomics Data Center (NGDC) with 492,167 strains in total (update to 2021/02/08). (B) Six O-glycosites in SARS-CoV-2 S protein are neither Ser nor Thr in SARS-CoV S protein, and were highlighted in red. The full-length sequence alignment was shown in Supplementary Figure 5. Sequences are derived from GenBank accession codes QHD43416 and AAP13441. See also Figure S4.

**Table 1.**
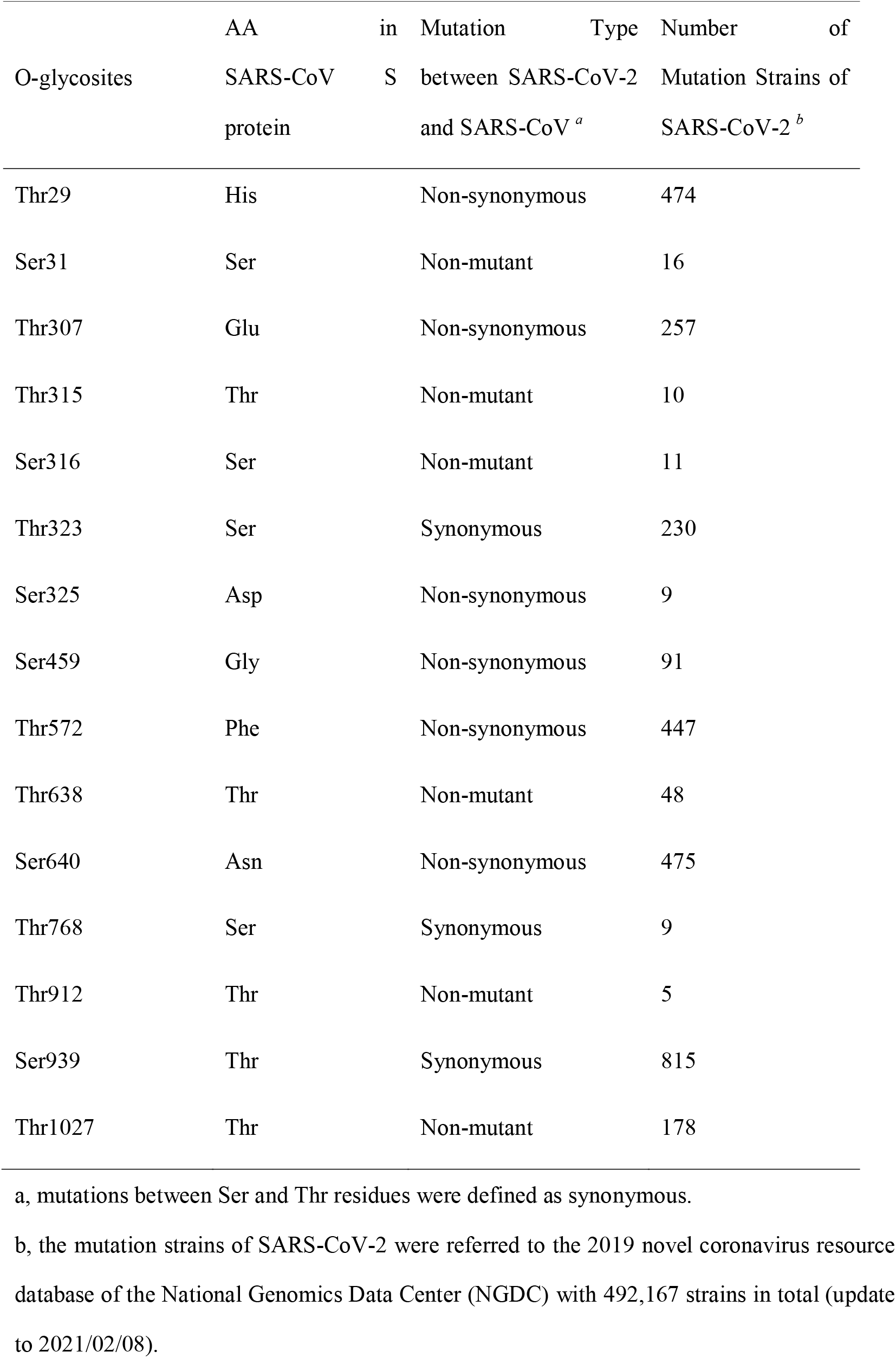
Summary of the mutations at O-glycosites of SARS-CoV-2 S protein.

## DISCUSSION

In this work, using MS method with HCD triggered ETD mode, together with peptide microarray assay and MD simulations, we reveal O-glycosites on SARS-CoV-2 S protein derived from HEK-293T cells and investigated how upregulated ppGalNAc-T6 expression might affect the S protein O-glycosylation and its trimeric stability. We identified a total of 15 O-glycosites and at least 10 different O-glycan structures on S protein. Importantly, ppGalNAc-T6 can dramatically change the O-glycosylation patterns of S protein in terms of the glycosites and glycan structures, supported by additional peptide microarray assays. Further MD simulations indicate that the interface O-glycosylation between adjacent promoters can potentially strengthen the S protein trimer structure via establishing nonpolar contacts or HB networks. Moreover, all the identified O-glycosites were highly conserved among currently discovered SRAS-CoV-2 strains. Our work provides deep insights into the molecular mechanism underlying the dynamic regulations of the S protein O-glycosylation during viral infection, exerted probably via upregulating certain O-glycosyltransferase in the host cell.

We employed an HCD triggered ETD MS approach to accurately pinpoint the glycosites and corresponding glycan structures on S protein. Using the HCD mode alone can potentially cause significant sugar neutral loss, thus the exact locations of the O-glycosites are hard to be undetermined(Vakhrushev et al., 2013). In contrast, the ETD mode can provide both glycosite and glycan-structure information owning to its capability of formatting the fragment ions of both glycan and peptide backbone (Riley et al., 2020). Here, by using the oxonium ion of sugar neutral loss in HCD to trigger ETD, and combining the spectra of HCD and ETD, we unambiguously determined all possible O-glycosites on S protein with a high degree of confidence(Wang et al., 2020).

Viral infections can modulate many host bioprocesses(Gordon et al., 2020; Shi et al., 2014). Recent studies revealed that SARS-CoV-2 could fine-tune the host cell functions including proteostasis, nucleotide biosynthesis and splicing(Bojkova et al., 2020; Messner et al., 2020). In the case of O-glycosylation processing, former studies have shown that the expression of host ppGalNAc-Ts could also be regulated by SARS-CoV-2 infections. Our site-specific O-glycosylation analysis revealed that the T6 overexpression would dramatically change the O-glycosylation patterns of S protein, which further help to stabilize its trimer structure since some of the O-glycosites locate within the promoter interface. Interesting, the O-glycosylation at two sites S316 and T323, were decreased upon T6 upregulation. Our peptide microarray assays reveal that the R319-to-P330 peptide tends to be glycosylated by T1 and T2 rather than T6, while the neighboring T307-to-F318 peptide prefers T6 glycosylation (Figure 2C). We thus speculate that the O-glycosylation of T307 and T315 by T6 might potentially impose an influence on the glycosylation of the neighboring S316 and T323 sites by T1 or T2, likely via steric hindrances.

Intriguingly, the T6 overexpression gives rise to increased O-glycosylation levels of T912, T678 and T572 that locate at the promoter interfaces, thereby strengthening the trimerization of S protein via forming stable interactions with adjacent promoters, as revealed by MD simulations. It is noteworthy that well-folded trimeric S protein was found to play essential roles in the SARS-CoV-2 infection. Thus, the SARS-CoV-2 infection might hijack the host T6 to dynamically upregulate the O-glycosylation level of S protein, thereby stabilizing the trimeric S protein structure and promoting the viral infection. Moreover, up to 20 ppGalNAc-Ts present in human(Peng et al., 2010), each has distinct expression patterns in different cells, tissues, or patients(Bennett et al., 2012). Therefore, SARS-CoV-2 from different individuals likely vary in the O-glycosylation fingerprints on the S protein, which might provide one possible reason for why the SARS-CoV-2 from different patients shows varied infectivity and antigenicity(Li et al., 2020a; Paggiaro et al., 2021).

## Acknowledgments

This work was supported by the National Science and Technology Major Project of China (2018ZX10302205), the National Natural Science Foundation of China (32071271, 31770850, and 22007065)

## Author Contributions

Y. Z. L.D and Z.X. conceived conception and design this study. Z.X. and J.S. prepared samples and Z.X. performed all experiments and data analysis. X.K. C.Z. and J.H. performed LC-MS/MS procedure. X.K., H.Z. and H.K. performed MS data analysis. Y.L. A.K. and S.T. checked peptide array data analysis, J.T. and L.D. performed molecular dynamics analysis. Z. X. X.K. and L.D wrote the original draft. W.Y., L.D., and Y.Z. reviewed manuscript and final approval of manuscript.

## EXPERIMENTAL MODEL AND SUBJECT DETAILS

### Cells

The Human Embryonic Kidney 293T (HEK-293T) cells were obtained from the Chinese National Human Genome Center. HEK-293T were cultured in High Glucose Dulbecco’s Modified Eagle Medium, supplemented with 10% heat-inactivated fetal bovine serum and incubated in a 5% CO2 atmosphere at 37 °C.

## METHOD DETAILS

### Materials

The recombinant SARS-CoV-2 S ectodomain (FLAG tag) were home-made and expressed in human embryonic kidney 293T (HEK-293T) cells. The SARS-CoV-2 S protein subunit 1 (His tag) expressed by HEK-293 cells were purchased from Sanyoubio (Shanghai, China). Dithiothreitol (DTT), iodoacetamide (IAA), formic acid (FA), trifluoroacetic acid (TFA), acetonitrile (ACN), ammonium bicarbonate (NH_4_HCO_3_), anti-FALG antibody, anti-FLAG M2 affinity gel, and neuraminidase were purchased from Sigma-Aldrich (St. Louis, MO, USA). Sequencing-grade trypsin were purchased from Enzyme & Spectrum (Beijing, China). C18 Tips were purchased from Thermo Fisher Scientific (Shanghai, China). The PNGase F, O-glycosidase and furin were purchased from NEB Labs (Shanghai, China).

### Protein expression and purification

To express the recombinant ectodomain of S protein (S-ECD), a mammalian-codon-optimized gene encoding SRAS-CoV-2 S (GenBank: QHD43416.1) residues 1−1208 with proline substitutions at residues 986 and 987,(Pallesen et al., 2017) a “GAAG” substitution at the furin cleavage site (residues 682−685) was synthesized and cloned into the mammalian expression vector pCMV-3×FLAG-14. This expression vector was used to transiently transfect HEK-293T cells using polyethylenimine. The C-terminal 3×FLAG tagged S-ECD protein (S-ECD-3×FLAG) was purified from cell medium or whole cell lysate using anti-FLAG M2 affinity gel, and then competitively eluted by 3×FLAG peptide.

### Glycosidase treatment

The purified S-ECD-3×FLAG proteins (50 ng) were treated with PNGaseF, neuraminidase and/or O-glycosidase according to the manufacturer’s protocol. Briefly, the proteins were denatured in 1× Glycoprotein Denaturing Buffer at 98°C for 10 min, followed by adding with GlycoBuffer 2 and NP-40. The proteins were then treated with PNGaseF, neuraminidase and/or O-glycosidase at 37°C for 2 h.

### Protein digestion

The N-glycans of S1 and S-ECD-3×FLAG proteins were removed by PNGaseF firstly, and then the proteins were separated using SDS-PAGE. The gel was stained by coomassie blue and the bands corresponding to S1 (70 to 110 kDa) and S-ECD-3×FLAG (130 to 170 kDa) were cut into ~1 mm^3^ cubes. Gel pieces were destained with 50% ACN in 50mM NH_4_HCO_3_ buffer at room temperature (RT) for 30 min, followed by dehydrating with 100% ACN. The proteins in gel were reduced by 10 mM DTT at 56°C for 45 min, followed by dehydrating with 100% ACN. The proteins in gel were then alkylated by 60 mM IAA at RT for 45 min in the dark. Subsequently, the proteins were digested by trypsin in 50 mM NH_4_HCO_3_) for 18 h at 37 °C. The peptides were extracted out from the gel by addition of 50% ACN containing 5% formic acid, and the released peptides were desalted using C18 Tips and speed-dried.

### Mass Spectrometry Analysis

LC-MS/MS analysis was performed by coupling a nanoLC (Dionex Ultimate 3000, Thermo Fisher Scientific) to an Orbitrap Fusion mass spectrometer (Thermo Fisher Scientific). Each sample was dissolved in 0.1% formic acid and was delivered to an analytical column (Dikma, inspire C18, 3 μm, Canada, 150 mm×75 μm, self-packed) and separated using a gradient at a flow rate of 0.3 μL/min: 0-3 min 1-4.8% B, 3-38 min 4.8-16% B; 37-47 min 16-28% B; 47-48 min 28-72% B; 48-60 min 72% B; 60-70 min 90% B, 70-80 min equilibration by 1% B (solvent A, 100% H_2_O; solvent B, 100% acetonitrile; both containing 0.1% (v/v) formic acid).

For tandem MS analysis, an “HCD triggered subsequent ETD scan” strategy was applied for most runs. Briefly, a precursor MS1 scan (m/z 300−1800) for intact peptides was acquired in the Orbitrap at 60,000 resolution, then followed by MS2 scans of the 20 most abundant multiply charged precursors in the MS1 spectrum (“top 20” method) using HCD fragmentation (m/z 100−2000, resolution 30,000) in Orbitrap mass analyzer. Subsequent ETD-based MS2 acquisition for the same precursor will be triggered (m/z 150−2000, resolution 30,000) and acquired in the Orbitrap when a HexNAc fragment at either m/z 204.0867, 186.0766 or 138.0545 was detected in the previous HCD-MS2 spectrum. The normalized collision energy for HCD was set to 30%. ETD fragmentation was performed with an automated calibrated charge-dependent reaction time supplemented by 15% HCD activation. Automatic gain control targets were 400,000 ions for MS1 and 50,000 ions for MS2 scans. Dynamic exclusion of 8 s was enabled to prevent repeated acquisition of the same precursor.

### Mass spectrometric data analysis

The LC-MS/MS raw data were analyzed manually for each glycopeptide using Xcalibur 3.0.63 (Thermo Fisher Scientific). The spectra were considered as O-glycopeptides only when the saccharide oxonium ions of HexNAc containing the pair of ions at m/z of 204.087 and 186.076 were detected. The O-glycosites and O-glycan structures were interpreted manually. The mass tolerances for precursors and fragment ions were set to 10 ppm and 0.02 Da, respectively. The glycosites and glycoforms of S protein purified from HEK-293T cells were confirmed according to the mass spectrometry results of commercial S1 using a ‘match between runs’ analysis. In brief, the same precursor ions (± 10 ppm) with a similar retention time (± 5 min) in the mass spectrometry results of commercial S1 or S protein purified from HEK-293T cells were considered as the same glycopeptides with same O-glycosites and O-glycans. The quantification of each glycan at each site was achieved by extracting the corresponding chromatographic areas. The areas of all charge states for a single glycopeptide were summed.

### On-chip ppGalNAc-Ts assay

The peptide microarray of S protein was home made as described previously(Li et al., 2020b). The on-chip ppGalNAc-Ts assay were modified from reported earlier(Xu et al., 2017). In brief, the microarrays were incubated with 5% w/v BSA/TBS (50 mM Tris and 150 mM NaCl, pH 7.4) for 1 h. The blocked microarrays were incubated overnight at 37□ with ppGalNAc-Ts reaction mixture containing 10 ng/μL ppGalNAc-Ts, 25 mM Tris-HCl (pH 7.4), 5 mM MnCl_2_, 0.2% v/v Triton X-100, and 0.5 mM UDP-GalNAc in 5% w/v BSA. The microarrays were then washed for 10 min three times in wash buffer (TBST containing 200 mM NaCl, 0.1% v/v Tween 20, and 0.2% w/v SDS), followed by the incubation with 1 μg/mL biotinylated VVA lectin (Vector Labs, CA, USA)1 h at RT. Then, the microarrays were washed for 10 min three times in wash buffer, followed by the incubation with 1 μg/mL Cy5 labeled streptavidin. Then, the microarrays were washed for 10 min three times in wash buffer. Microarrays were scanned at Ex 532 by a GenePix 4200A slide scanner (Molecular Devices, CA, USA). The foreground intensities and background intensities were extracted by the GenePix Pro 6.0 software from the microarray images. The S/N of a protein was averaged from triplicate spots.

### 3D modeling and molecular dynamics simulations

For each of the six protomer-interface O-glycosites, including T1027, T912, T768, S316, T315, and T572, we constructed the Tn-modified S-protein based on one crystal structure of S-protein trimer (pdb ID: 6xr8)(Cai et al., 2020). We then solvated each O-glycosylated S-trimer in a cubic box filled with TIP3P water. 3 Na^+^ ions were added to neutralize the whole system. The final system was then subjected to energy minimization to relieve local steric clashes, followed by a heating simulation from 0 to 310 K within 50 ps. Next, we performed 500 ps equilibrium MD simulations at 310 K by constraining all solute heavy atoms. Finally, we conducted 20-ns molecular dynamics (MD) simulations for each system by keeping the temperature at 310 K, controlled by the Langevin thermostat(Wu and Brooks, 2003). All the MD simulations were performed using the Amber14 package(Case et al., 2014). The ff14SB and GLYCAM06j-1 force fields were employed to describe the protein and GalNAc, respectively(Kirschner et al., 2008; Maier et al., 2015). The SHAKE algorithm was used to constrain the bond lengths involving hydrogen atoms(Ryckaert et al., 1977). The non-bonded cutoff distance was set as 10 Å, and the long-range electrostatic interaction was calculated using the particle mesh Ewald (PME) method(Essmann et al., 1995). As a control, we also performed one 20-ns MD simulations for the un-glycoslated S-protein trimer.

### Supplemental Table

Table S1. Detail LC-MS/MS information of all the O-glycopeptides detected in the three samples of S-ECD, S-ECD co-expressed with T6, and S1. (Related to Figure 1, 3 and S2)

## References

Bennett, E.P., Mandel, U., Clausen, H., Gerken, T.A., Fritz, T.A., and Tabak, L.A. (2012). Control of mucin-type O-glycosylation: a classification of the polypeptide GalNAc-transferase gene family. Glycobiology 22, 736–756. doi:10.1093/glycob/cwr182.

Bojkova, D., Klann, K., Koch, B., Widera, M., Krause, D., Ciesek, S., Cinatl, J., and Munch, C. (2020). Proteomics of SARS-CoV-2-infected host cells reveals therapy targets. Nature 583, 469–472. doi:10.1038/s41586-020-2332-7.

Cai, Y., Zhang, J., Xiao, T., Peng, H., Sterling, S.M., Walsh, R.M., Jr., Rawson, S., Rits-Volloch, S., and Chen, B. (2020). Distinct conformational states of SARS-CoV-2 spike protein. Science 369, 1586–1592. doi:10.1126/science.abd4251.

Casalino, L., Gaieb, Z., Goldsmith, J.A., Hjorth, C.K., Dommer, A.C., Harbison, A.M., Fogarty, C.A., Barros, E.P., Taylor, B.C., McLellan, J.S., et al. (2020). Beyond Shielding: The Roles of Glycans in the SARS-CoV-2 Spike Protein. ACS Cent Sci 6, 1722–1734. doi:10.1021/acscentsci.0c01056.

Case, D.A., Babin, V., Berryman, J., Betz, R.M., Cai, Q., Cerutti, D.S., Cheatham III, T.E., Darden, T.A., Duke, R.E., Gohlke, H., et al. (2014). AMBER 14 (University of California, San Francisco, http://ambermd.org/.).

Chan, J.F., Kok, K.H., Zhu, Z., Chu, H., To, K.K., Yuan, S., and Yuen, K.Y. (2020a). Genomic characterization of the 2019 novel human-pathogenic coronavirus isolated from a patient with atypical pneumonia after visiting Wuhan. Emerg Microbes Infect 9, 221–236. doi:10.1080/22221751.2020.1719902.

Chan, J.F., Yuan, S., Kok, K.H., To, K.K., Chu, H., Yang, J., Xing, F., Liu, J., Yip, C.C., Poon, R.W., et al. (2020b). A familial cluster of pneumonia associated with the 2019 novel coronavirus indicating person-to-person transmission: a study of a family cluster. Lancet 395, 514–523. doi:10.1016/S0140-6736(20)30154-9.

Chen, N., Zhou, M., Dong, X., Qu, J., Gong, F., Han, Y., Qiu, Y., Wang, J., Liu, Y., Wei, Y., et al. (2020). Epidemiological and clinical characteristics of 99 cases of 2019 novel coronavirus pneumonia in Wuhan, China: a descriptive study. Lancet 395, 507–513. doi:10.1016/S0140-6736(20)30211-7.

De Las Rivas, M., Paul Daniel, E.J., Narimatsu, Y., Companon, I., Kato, K., Hermosilla, P., Thureau, A., Ceballos-Laita, L., Coelho, H., Bernado, P., et al. (2020). Molecular basis for fibroblast growth factor 23 O-glycosylation by GalNAc-T3. Nat Chem Biol 16, 351–360. doi:10.1038/s41589-019-0444-x.

Essmann, U., Perera, L., Berkowitz, M.L., Darden, T., Lee, H., and Pedersen, L.G. (1995). A smooth particle mesh Ewald method. J Chem Phys 103, 8577–8593. doi:10.1063/1.470117

Gao, C., Zeng, J., Jia, N., Stavenhagen, K., Matsumoto, Y., Zhang, H., Li, J., Hume, A.J., Muhlberger, E., van Die, I., et al. (2020). SARS-CoV-2 Spike Protein Interacts with Multiple Innate Immune Receptors. bioRxiv. doi:10.1101/2020.07.29.227462.

Gerken, T.A., Jamison, O., Perrine, C.L., Collette, J.C., Moinova, H., Ravi, L., Markowitz, S.D., Shen, W., Patel, H., and Tabak, L.A. (2011). Emerging paradigms for the initiation of mucin-type protein O-glycosylation by the polypeptide GalNAc transferase family of glycosyltransferases. J Biol Chem 286, 14493–14507. doi:10.1074/jbc.M111.218701.

Gong, Z., Zhu, J.W., Li, C.P., Jiang, S., Ma, L.N., Tang, B.X., Zou, D., Chen, M.L., Sun, Y.B., Song, S.H., et al. (2020). An online coronavirus analysis platform from the National Genomics Data Center. Zool Res 41, 705–708. doi:10.24272/j.issn.2095-8137.2020.065.

Gordon, D.E., Jang, G.M., Bouhaddou, M., Xu, J.W., Obernier, K., White, K.M., O’Meara, M.J., Rezelj, V.V., Guo, J.F.Z., Swaney, D.L., et al. (2020). A SARS-CoV-2 protein interaction map reveals targets for drug repurposing. Nature 583, 459–468. doi:10.1038/s41586-020-2286-9.

Goth, C.K., Vakhrushev, S.Y., Joshi, H.J., Clausen, H., and Schjoldager, K.T. (2018). Fine-Tuning Limited Proteolysis: A Major Role for Regulated Site-Specific O-Glycosylation. Trends Biochem Sci 43, 269–284. doi:10.1016/j.tibs.2018.02.005.

Hao, Y., Fan, X., Shi, Y., Zhang, C., Sun, D.E., Qin, K., Qin, W., Zhou, W., and Chen, X. (2019). Next-generation unnatural monosaccharides reveal that ESRRB O-GlcNAcylation regulates pluripotency of mouse embryonic stem cells. Nat Commun 10, 4065. doi:10.1038/s41467-019-11942-y.

Hoffmann, M., Kleine-Weber, H., Schroeder, S., Kruger, N., Herrler, T., Erichsen, S., Schiergens, T.S., Herrler, G., Wu, N.H., Nitsche, A., et al. (2020). SARS-CoV-2 Cell Entry Depends on ACE2 and TMPRSS2 and Is Blocked by a Clinically Proven Protease Inhibitor. Cell 181, 271–280. doi:10.1016/j.cell.2020.02.052.

Kirschner, K.N., Yongye, A.B., Tschampel, S.M., González-Outeiriño, J., Daniels, C.R., Foley, B.L., and Woods, R.J. (2008). GLYCAM06: A generalizable biomolecular force field. Carbohydrates. J Comput Chem 29, 622–655. doi:10.1002/jcc.20820.

Kong, Y., Joshi, H.J., Schjoldager, K.T., Madsen, T.D., Gerken, T.A., Vester-Christensen, M.B., Wandall, H.H., Bennett, E.P., Levery, S.B., Vakhrushev, S.Y., et al. (2015). Probing polypeptide GalNAc-transferase isoform substrate specificities by in vitro analysis. Glycobiology 25, 55–65. doi:10.1093/glycob/cwu089.

Kudelka, M.R., Ju, T., Heimburg-Molinaro, J., and Cummings, R.D. (2015). Simple sugars to complex disease--mucin-type O-glycans in cancer. Adv Cancer Res 126, 53–135. doi:10.1016/bs.acr.2014.11.002.

Lan, J., Ge, J., Yu, J., Shan, S., Zhou, H., Fan, S., Zhang, Q., Shi, X., Wang, Q., Zhang, L., et al. (2020). Structure of the SARS-CoV-2 spike receptor-binding domain bound to the ACE2 receptor. Nature 581, 215–220. doi:10.1038/s41586-020-2180-5.

Lermyte, F., Valkenborg, D., Loo, J.A., and Sobott, F. (2018). Radical solutions: Principles and application of electron-based dissociation in mass spectrometry-based analysis of protein structure. Mass Spectrom Rev 37, 750–771. doi:10.1002/mas.21560.

Li, Q., Wu, J., Nie, J., Zhang, L., Hao, H., Liu, S., Zhao, C., Zhang, Q., Liu, H., Nie, L., et al. (2020a). The Impact of Mutations in SARS-CoV-2 Spike on Viral Infectivity and Antigenicity. Cell 182, 1284–1294 e1289. doi:10.1016/j.cell.2020.07.012.

Li, X., Wang, J., Li, W., Xu, Y., Shao, D., Xie, Y., Xie, W., Kubota, T., Narimatsu, H., and Zhang, Y. (2012). Characterization of ppGalNAc-T18, a member of the vertebrate-specific Y subfamily of UDP-N-acetyl-alpha-D-galactosamine: polypeptide N-acetylgalactosaminyltransferases. Glycobiology 22, 602–615. doi:10.1093/glycob/cwr179.

Li, Y., Lai, D.Y., Zhang, H.N., Jiang, H.W., Tian, X., Ma, M.L., Qi, H., Meng, Q.F., Guo, S.J., Wu, Y., et al. (2020b). Linear epitopes of SARS-CoV-2 spike protein elicit neutralizing antibodies in COVID-19 patients. Cell Mol Immunol 17, 1095–1097. doi:10.1038/s41423-020-00523-5.

Liao, M., Liu, Y., Yuan, J., Wen, Y., Xu, G., Zhao, J., Cheng, L., Li, J., Wang, X., Wang, F., et al. (2020). Single-cell landscape of bronchoalveolar immune cells in patients with COVID-19. Nat Med 26, 842–844. doi:10.1038/s41591-020-0901-9.

Liu, F., Xu, K., Xu, Z., de Las Rivas, M., Wang, C., Li, X., Lu, J., Zhou, Y., Delso, I., Merino, P., et al. (2017). The small molecule luteolin inhibits N-acetyl-alpha-galactosaminyltransferases and reduces mucin-type O-glycosylation of amyloid precursor protein. J Biol Chem 292, 21304–21319. doi:10.1074/jbc.M117.814202.

Maier, J.A., Martinez, C., Kasavajhala, K., Wickstrom, L., Hauser, K.E., and Simmerling, C. (2015). ff14SB: Improving the Accuracy of Protein Side Chain and Backbone Parameters from ff99SB. J Chem Theory Comput 11, 3696–3713. doi:10.1021/acs.jctc.5b00255.

Messner, C.B., Demichev, V., Wendisch, D., Michalick, L., White, M., Freiwald, A., Textoris-Taube, K., Vernardis, S.I., Egger, A.S., Kreidl, M., et al. (2020). Ultra-High-Throughput Clinical Proteomics Reveals Classifiers of COVID-19 Infection. Cell Syst 11, 11–24. doi:10.1016/j.cels.2020.05.012.

Paggiaro, A.O., Carvalho, V.F., and Gemperli, R. (2021). Effect of different human tissue processing techniques on SARS-CoV-2 inactivation-review. Cell Tissue Bank 22, 1–10. doi:10.1007/s10561-020-09869-6.

Pallesen, J., Wang, N.S., Corbett, K.S., Wrapp, D., Kirchdoerfer, R.N., Turner, H.L., Cottrell, C.A., Becker, M.M., Wang, L.S., Shi, W., et al. (2017). Immunogenicity and structures of a rationally designed prefusion MERS-CoV spike antigen. Proc Natl Acad Sci U S A 114, E7348–E7357. doi:10.1073/pnas.1707304114.

Peng, C., Togayachi, A., Kwon, Y.D., Xie, C., Wu, G., Zou, X., Sato, T., Ito, H., Tachibana, K., Kubota, T., et al. (2010). Identification of a novel human UDP-GalNAc transferase with unique catalytic activity and expression profile. Biochem Biophys Res Commun 402, 680–686. doi:10.1016/j.bbrc.2010.10.084.

Riley, N.M., Malaker, S.A., Driessen, M.D., and Bertozzi, C.R. (2020). Optimal Dissociation Methods Differ for N- and O-Glycopeptides. J Proteome Res 19, 3286–3301. doi:10.1021/acs.jproteome.0c00218.

Ryckaert, J.-P., Ciccotti, G., and Berendsen, H.J.C. (1977). Numerical integration of the cartesian equations of motion of a system with constraints: molecular dynamics of n-alkanes. J Comput Phys 23, 327–341. doi:10.1016/0021-9991(77)90098-5.

Sanda, M., Morrison, L., and Goldman, R. (2021). N- and O-Glycosylation of the SARS-CoV-2 Spike Protein. Anal Chem 93, 2003–2009. doi:10.1021/acs.analchem.0c03173.

Schjoldager, K.T., Joshi, H.J., Kong, Y., Goth, C.K., King, S.L., Wandall, H.H., Bennett, E.P., Vakhrushev, S.Y., and Clausen, H. (2015). Deconstruction of O-glycosylation--GalNAc-T isoforms direct distinct subsets of the O-glycoproteome. EMBO Rep 16, 1713–1722. doi:10.15252/embr.201540796.

Shajahan, A., Supekar, N.T., Gleinich, A.S., and Azadi, P. (2020). Deducing the N- and O-glycosylation profile of the spike protein of novel coronavirus SARS-CoV-2. Glycobiology 30, 981–988. doi:10.1093/glycob/cwaa042.

Shi, C.S., Qi, H.Y., Boularan, C., Huang, N.N., Abu-Asab, M., Shelhamer, J.H., and Kehrl, J.H. (2014). SARS-Coronavirus Open Reading Frame-9b Suppresses Innate Immunity by Targeting Mitochondria and the MAVS/TRAF3/TRAF6 Signalosome. J Immunol 193, 3080–3089. doi:10.4049/jimmunol.1303196.

Song, S., Ma, L., Zou, D., Tian, D., Li, C., Zhu, J., Chen, M., Wang, A., Ma, Y., Li, M., et al. (2020). The Global Landscape of SARS-CoV-2 Genomes, Variants, and Haplotypes in 2019nCoVR. Genomics Proteomics Bioinformatics. doi:10.1016/j.gpb.2020.09.001.

Steentoft, C., Vakhrushev, S.Y., Joshi, H.J., Kong, Y., Vester-Christensen, M.B., Schjoldager, K.T., Lavrsen, K., Dabelsteen, S., Pedersen, N.B., Marcos-Silva, L., et al. (2013). Precision mapping of the human O-GalNAc glycoproteome through SimpleCell technology. EMBO J 32, 1478–1488. doi:10.1038/emboj.2013.79.

Steentoft, C., Vakhrushev, S.Y., Vester-Christensen, M.B., Schjoldager, K.T., Kong, Y., Bennett, E.P., Mandel, U., Wandall, H., Levery, S.B., and Clausen, H. (2011). Mining the O-glycoproteome using zinc-finger nuclease-glycoengineered SimpleCell lines. Nat Methods 8, 977–982. doi:10.1038/nmeth.1731.

Stone, J.A., Nicola, A.V., Baum, L.G., and Aguilar, H.C. (2016). Multiple Novel Functions of Henipavirus O-glycans: The First O-glycan Functions Identified in the Paramyxovirus Family. PLoS Pathog 12, e1005445. doi:10.1371/journal.ppat.1005445.

Stukalov, A., Girault, V., Grass, V., Bergant, V., Karayel, O., Urban, C., Haas, D., Huang, Y., Oubraham, L., Wang, A., et al. (2020). Multi-level proteomics reveals host-perturbation strategies of SARS-CoV-2 and SARS-CoV. bioRxiv. doi:10.1101/2020.06.17.156455

Vakhrushev, S.Y., Steentoft, C., Vester-Christensen, M.B., Bennett, E.P., Clausen, H., and Levery, S.B. (2013). Enhanced mass spectrometric mapping of the human GalNAc-type O-glycoproteome with SimpleCells. Mol Cell Proteomics 12, 932–944. doi:10.1074/mcp.O112.021972.

Wang, D., Baudys, J., Bundy, J.L., Solano, M., Keppel, T., and Barr, J.R. (2020). Comprehensive Analysis of the Glycan Complement of SARS-CoV-2 Spike Proteins Using Signature Ions-Triggered Electron-Transfer/Higher-Energy Collisional Dissociation (EThcD) Mass Spectrometry. Anal Chem 92, 14730–14739. doi:10.1021/acs.analchem.0c03301.

Wang, S.J., Mao, Y., Narimatsu, Y., Ye, Z.L., Tian, W.H., Goth, C.K., Lira-Navarrete, E., Pedersen, N.B., Benito-Vicente, A., Martin, C., et al. (2018). Site-specific O-glycosylation of members of the low-density lipoprotein receptor superfamily enhances ligand interactions. J Biol Chem 293, 7408–7422. doi:10.1074/jbc.M117.817981.

Watanabe, Y., Allen, J.D., Wrapp, D., McLellan, J.S., and Crispin, M. (2020). Site-specific glycan analysis of the SARS-CoV-2 spike. Science 369, 330–333. doi:10.1126/science.abb9983.

Wu, X., and Brooks, B.R. (2003). Self-guided Langevin dynamics simulation method. Chem Phys Lett 381, 512–518. doi:10.1016/j.cplett.2003.10.013.

Xu, Z., Li, X., Zhou, S., Xie, W., Wang, J., Cheng, L., Wang, S., Guo, S., Cao, X., Zhang, M., et al. (2017). Systematic identification of the protein substrates of UDP-GalNAc:polypeptide N-acetylgalactosaminyltransferase-T1/T2/T3 using a human proteome microarray. Proteomics 17. doi:10.1002/pmic.201600485.

Yan, R., Zhang, Y., Li, Y., Xia, L., Guo, Y., and Zhou, Q. (2020). Structural basis for the recognition of SARS-CoV-2 by full-length human ACE2. Science 367, 1444–1448. doi:10.1126/science.abb2762.

Zhang, Y., Zhao, W., Mao, Y., Chen, Y., Zhu, J., Hu, L., Gong, M., Cheng, J., and Yang, H. (2020). Mucin-type O-glycosylation Landscapes of SARS-CoV-2 Spike Proteins. bioRxiv. doi:10.1101/2020.07.29.227785.

Zhao, P., Praissman, J.L., Grant, O.C., Cai, Y., Xiao, T., Rosenbalm, K.E., Aoki, K., Kellman, B.P., Bridger, R., Barouch, D.H., et al. (2020a). Virus-Receptor Interactions of Glycosylated SARS-CoV-2 Spike and Human ACE2 Receptor. Cell Host Microbe 28, 586–601 e586. doi:10.1016/j.chom.2020.08.004.

Zhao, W.M., Song, S.H., Chen, M.L., Zou, D., Ma, L.N., Ma, Y.K., Li, R.J., Hao, L.L., Li, C.P., Tian, D.M., et al. (2020b). The 2019 novel coronavirus resource. Yi Chuan 42, 212–221. doi:10.16288/j.yczz.20-030.

