## Supplemental information for "Altered O-glycosylation Level of SARS-CoV-2 Spike Protein by Host O-glycosyltransferase Strengthens Its Trimeric Structure"

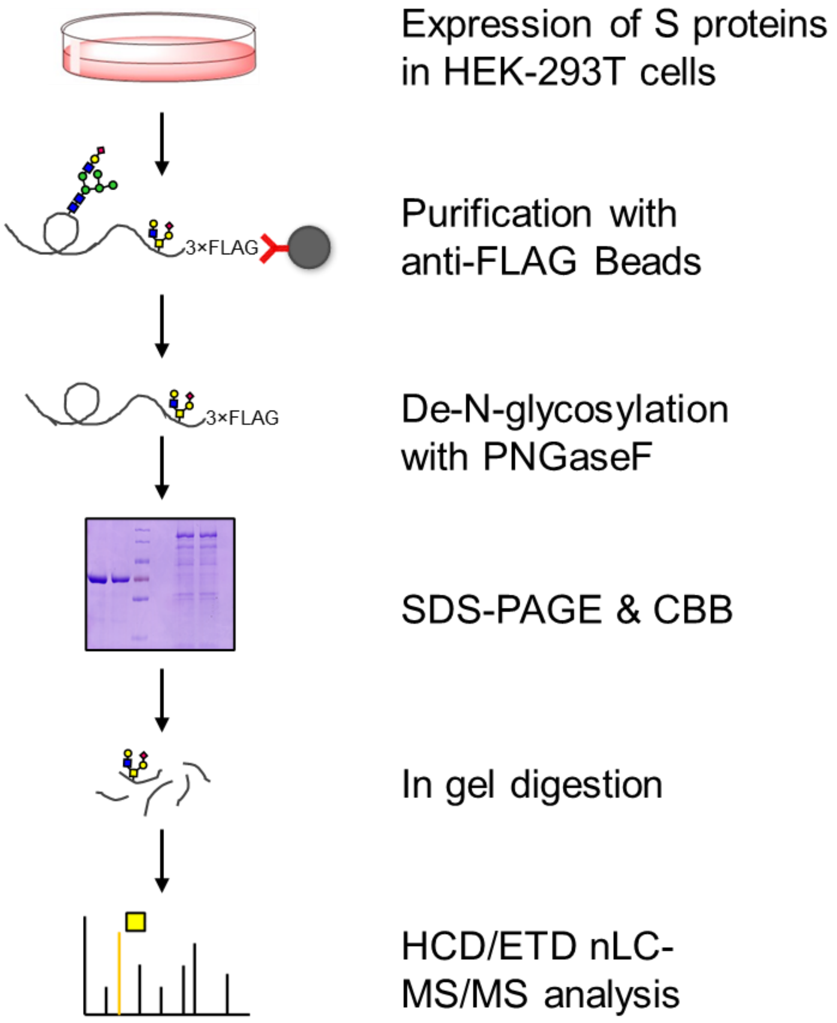


**Figure S1. Scheme of O-glycosylation analysis of SARS-CoV-2 S proteins.**

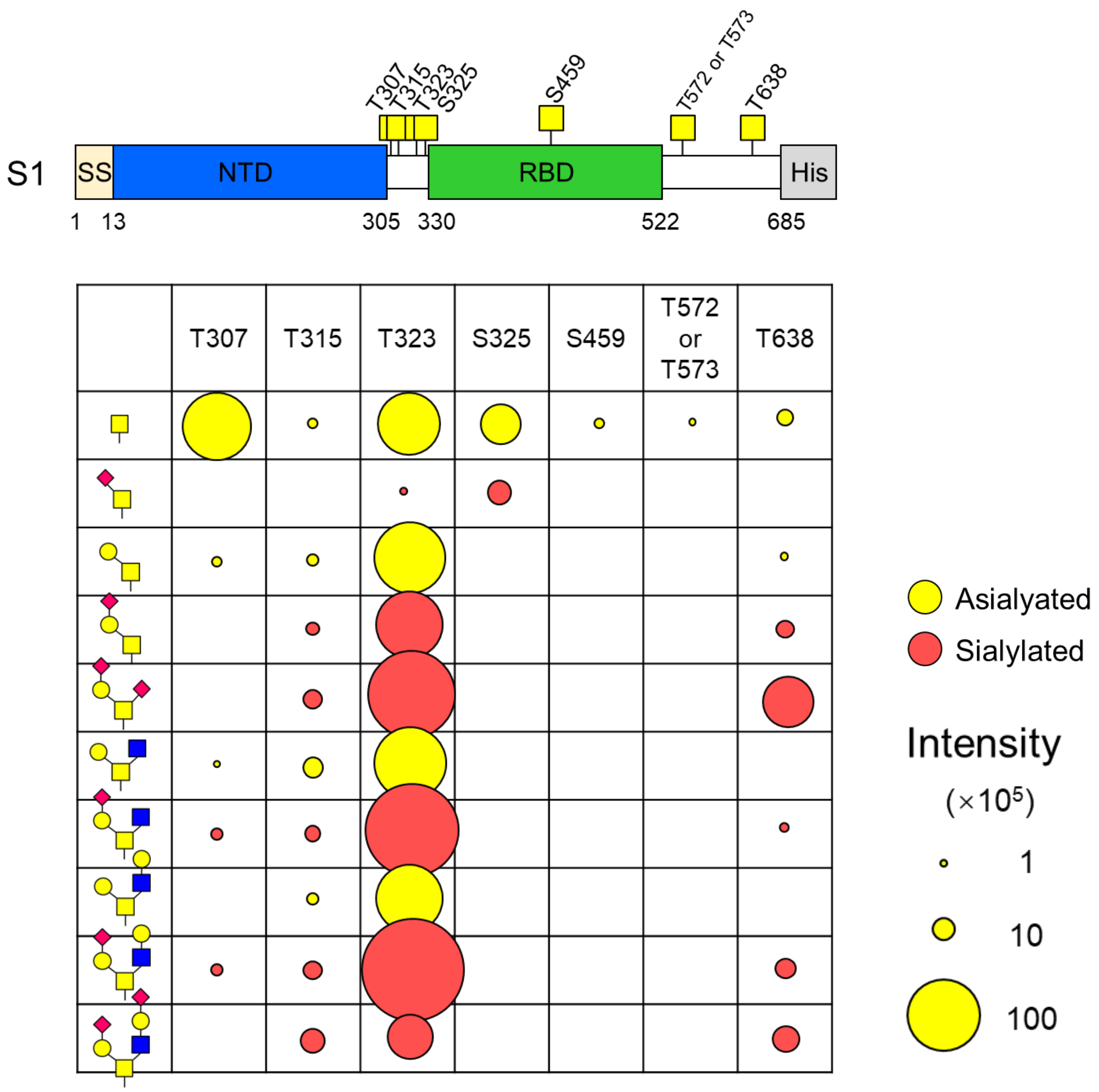


**Figure S2. Identification of the O-glycosites and O-glycans on S protein subunit 1.** Bubble areas indicate the relative intensities of each O-glycans at individual sites. Red bubbles indicate sialylated O-glycans, yellow bubbles indicate asialylated O-glycans.

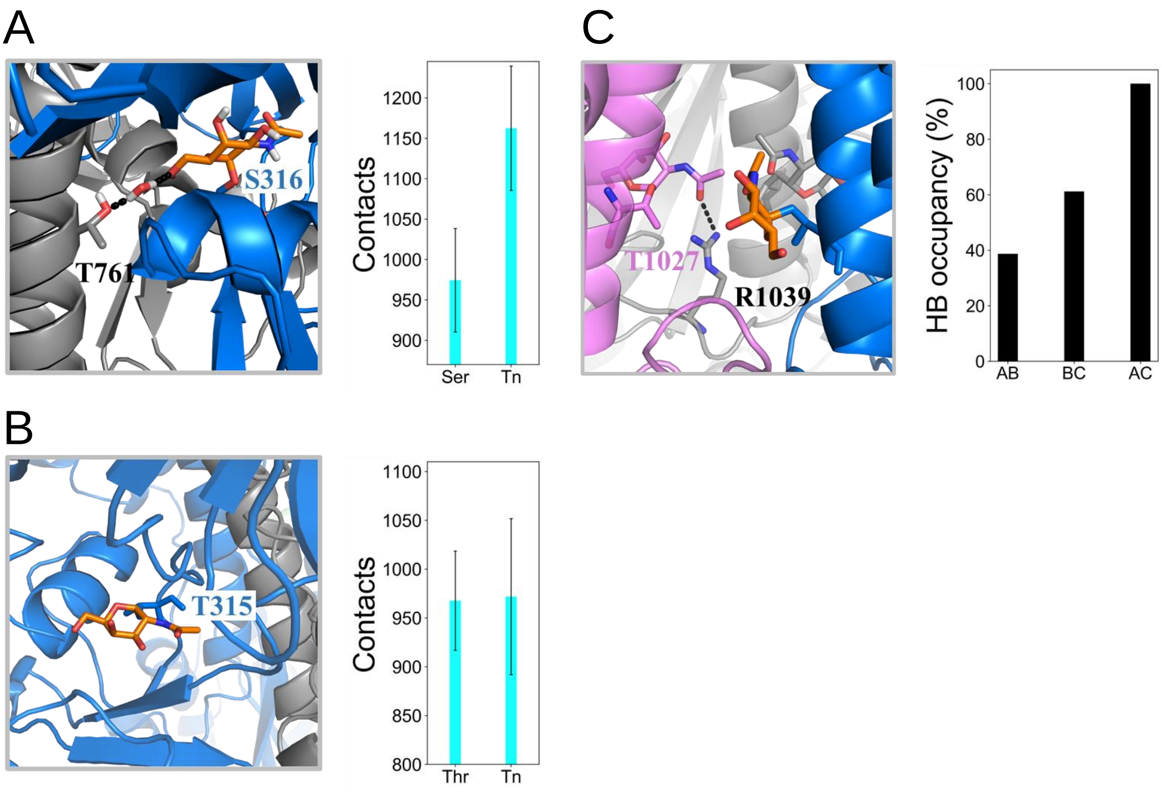


**Figure S3.** (A) The structural diagram with S316 glycosylated with GalNAc (Left panel); the contact numbers between each two neighbor protomers (Right panel); (B) The structural diagram with T315 glycosylated with GalNAc (Left panel); the contact numbers between each two neighbor protomers (Right panel); (C) The structural diagram with T1027 glycosylated with GalNAc (Left panel); the occupancy of HB formed by the GalNAc and R1039 from the neighbour protomer (Right panel)

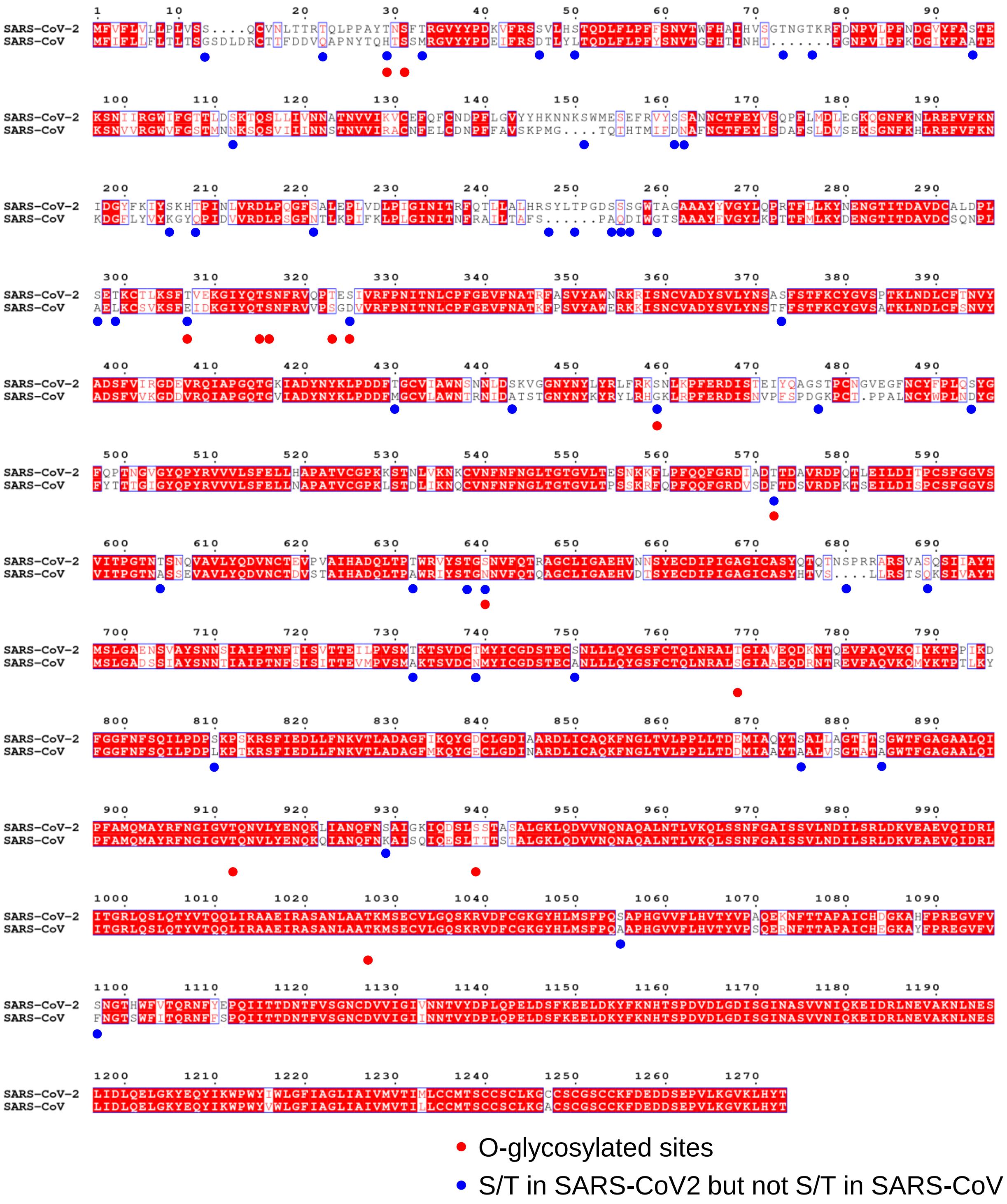


**Figure S4. Sequence alignment of SARS-CoV-2 S protein and SARS-CoV S protein.** Sequences shown are from GenBank accession codes QHD43416 and AAP13441. The O-glycosylated sites of SARS-CoV-2 S protein were labeled out with red spots. Forty-eight S/T residues in SARS-CoV2 S protein were not S/T amino acids in SARS-CoV S protein, and were labeled out with blue spots.

**Figure S5 Representative HCD and ETD MS/MS spectra. (Related to Figure 1, 3 and S2)**

A. Representative HCD and ETD MS/MS spectra of intact O-glycopeptides of S protein subunit 1 with determined O-glycosites.

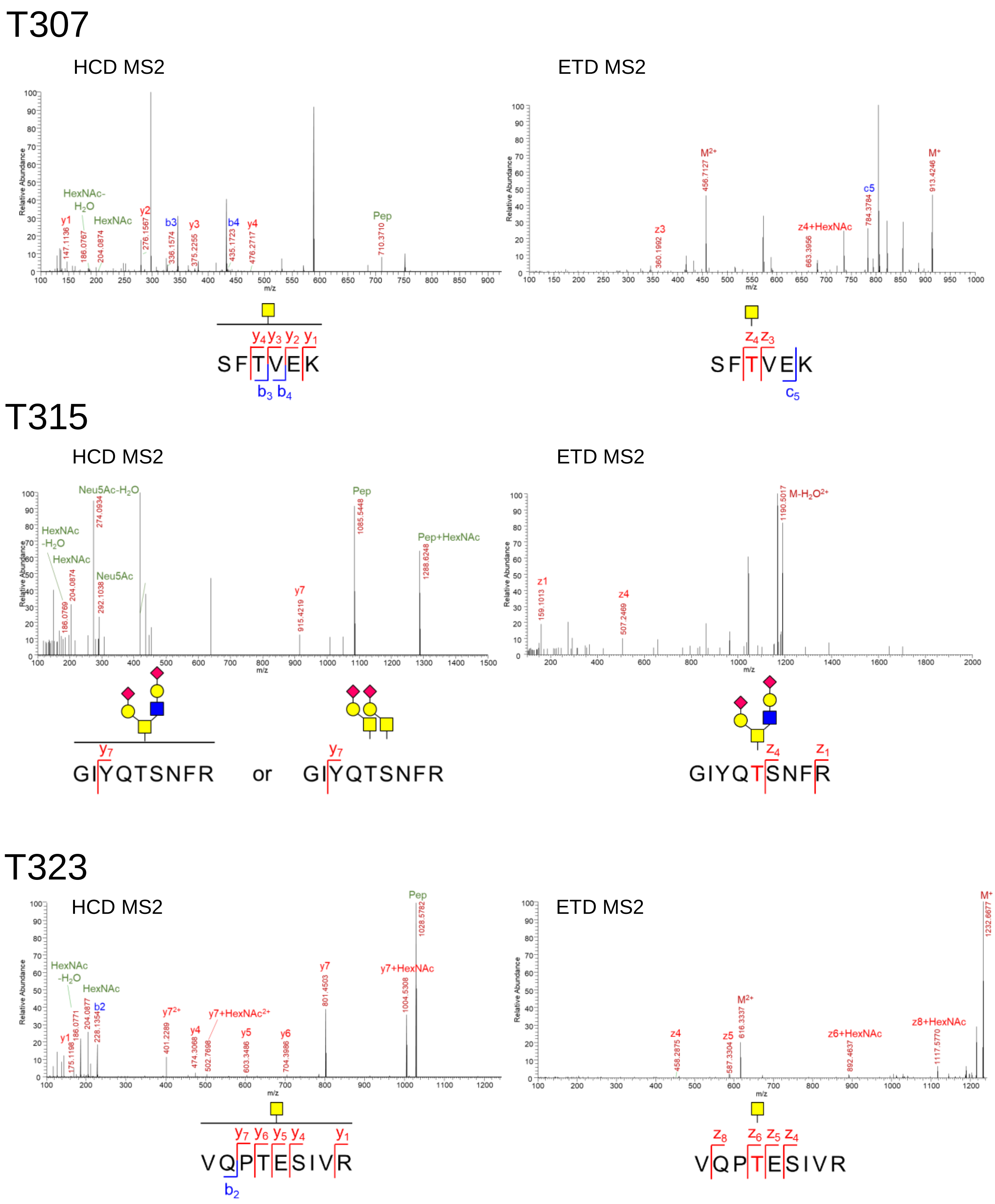

# T325 & S325

HCD MS2

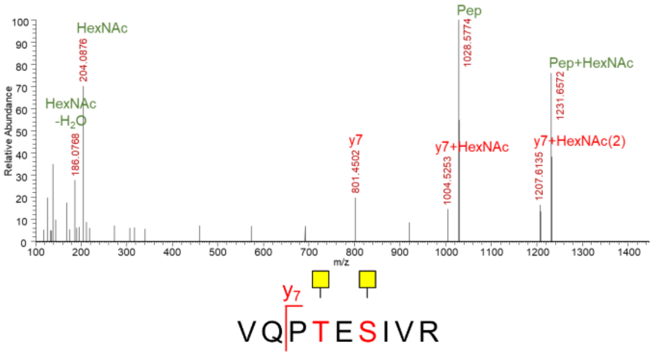


ETD MS2

Not detected

# S459

HCD MS2

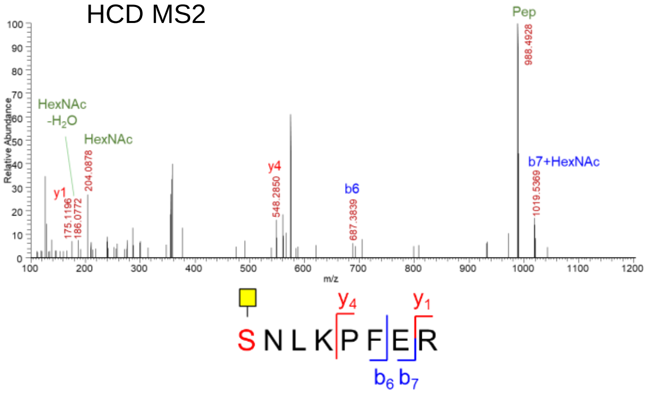


ETD MS2

Not detected

# T638

HCD MS2

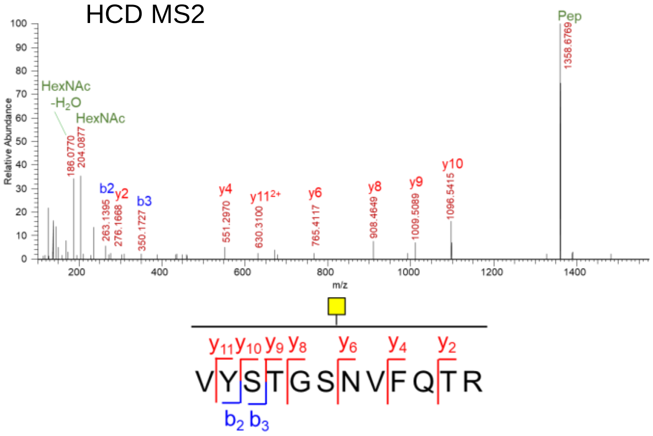


ETD MS2

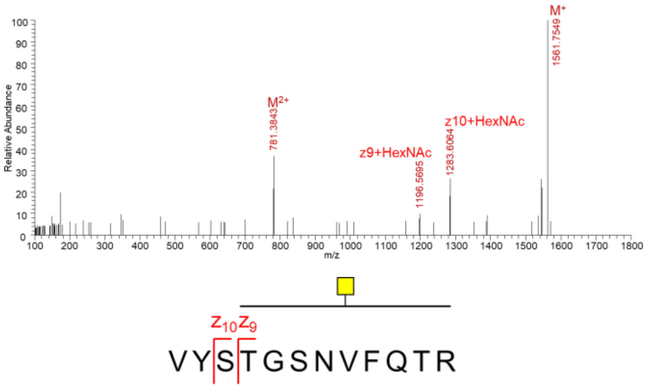


This site was confirmed in the MS/MS spectrum of S-ECD-3×FLAG protein in Fig. S5.

**Figure S5 Representative HCD and ETD MS/MS spectra. (Related to Figure 1, 3 and S2)**

B. Representative HCD and ETD MS/MS spectra of intact O-glycopeptides with all O-glycans at T323 of S1 subunit.

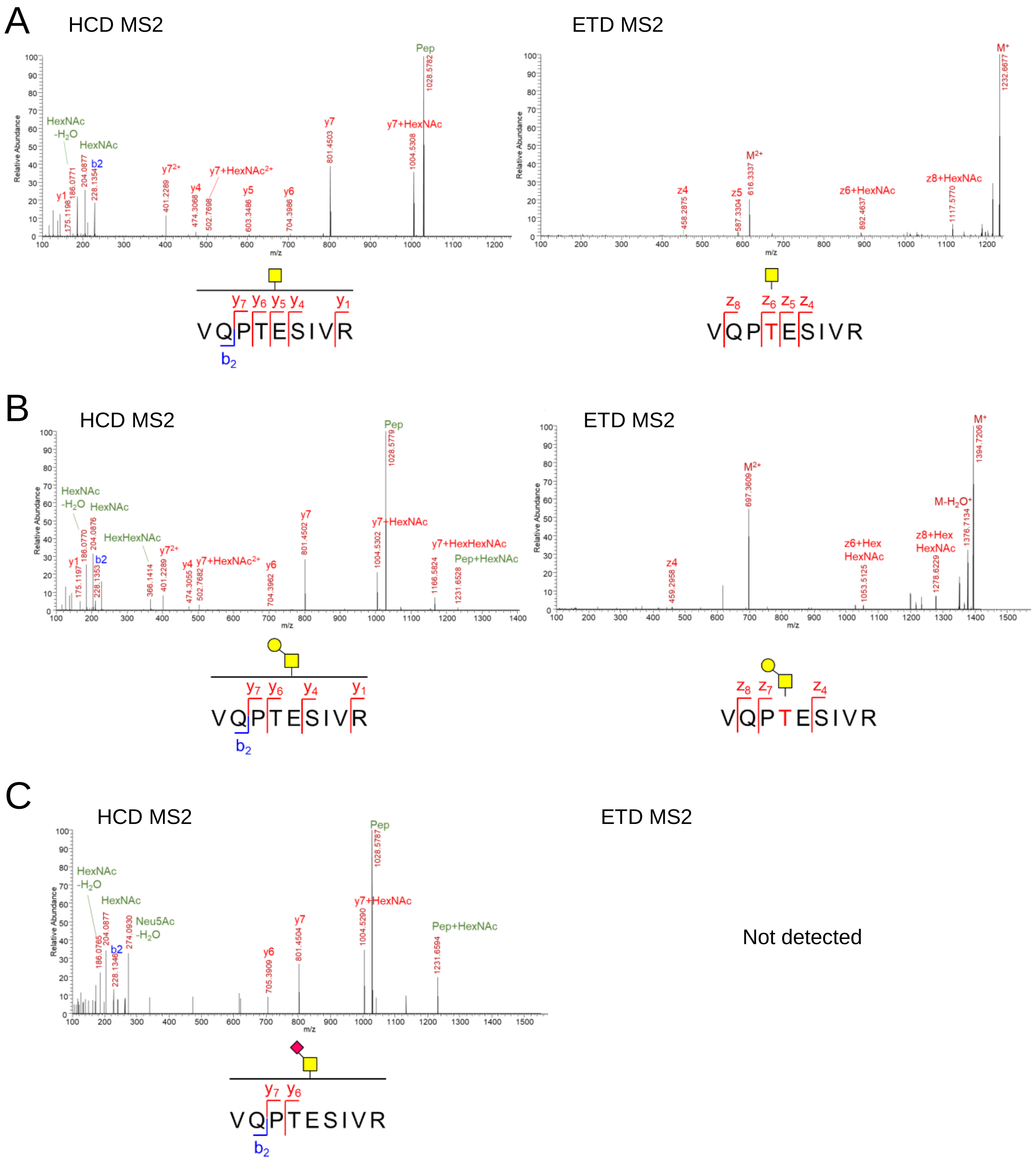

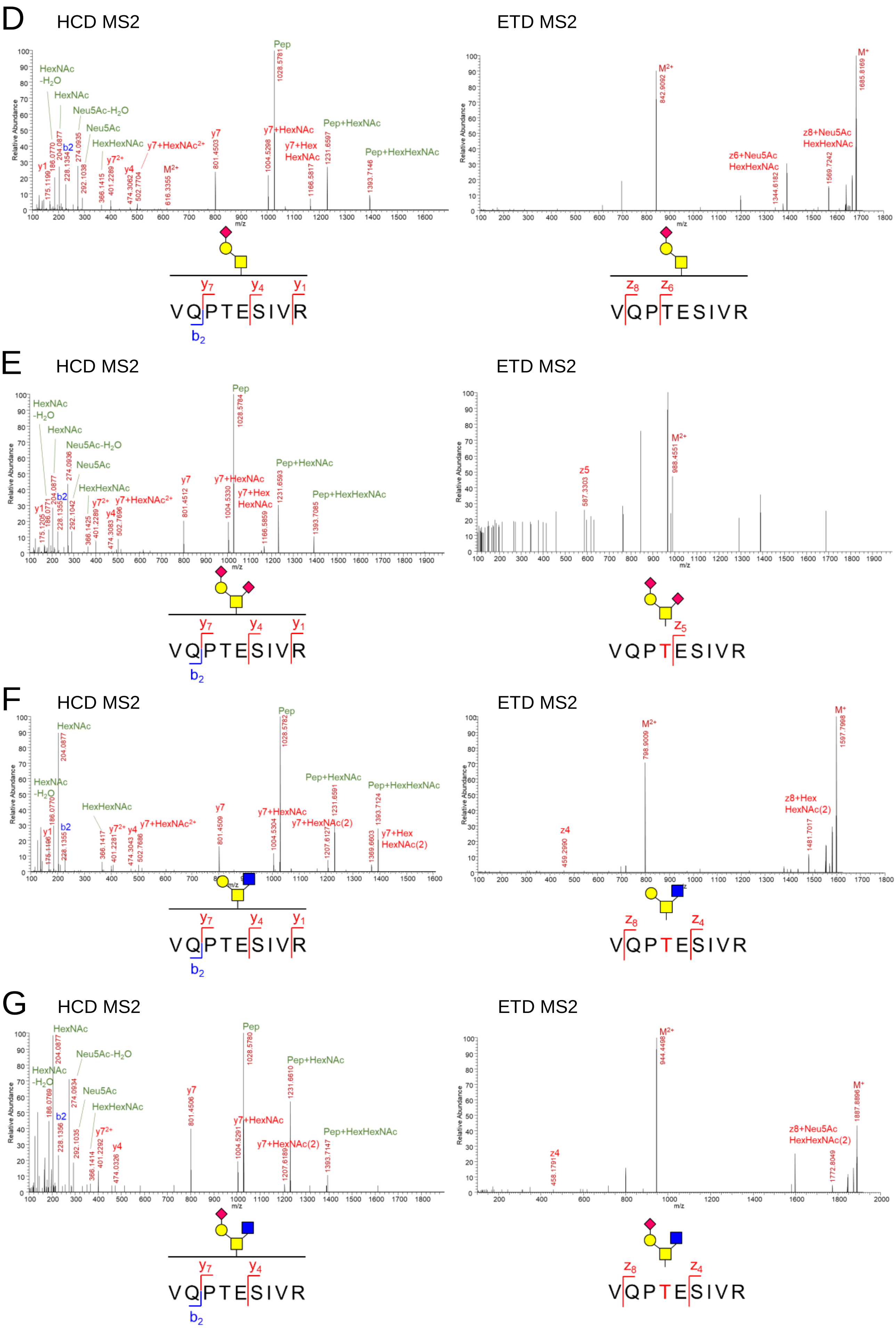

D
HCD MS2
ETD MS2
Pep
HexNAc -H2O
HexNAc
Neu5Ac-H2O
Neu5Ac
HexHexNAc
y7+HexNAc2+
y7
y7+HexNAc
y7+Hex HexNAc
Pep+HexNAc
Pep+HexHexNAc
y1
b2
y72+
y4
M2+
M+
z8+Neu5Ac HexHexNAc
z6+Neu5Ac HexHexNAc
Relative Abundance
m/z
y7 y4 y1
VQPTESIVR
b2
z8 z6
VQPTESIVR
E
HCD MS2
ETD MS2
z5
VQPTESIVR
F
HCD MS2
ETD MS2
y7+HexNAc(2)
y7+Hex HexNAc(2)
z8+Hex HexNAc(2)
z4
z8 z4
VQPTESIVR
G
HCD MS2
ETD MS2
z8+Neu5Ac HexHexNAc(2)
z8 z4
VQPTESIVR

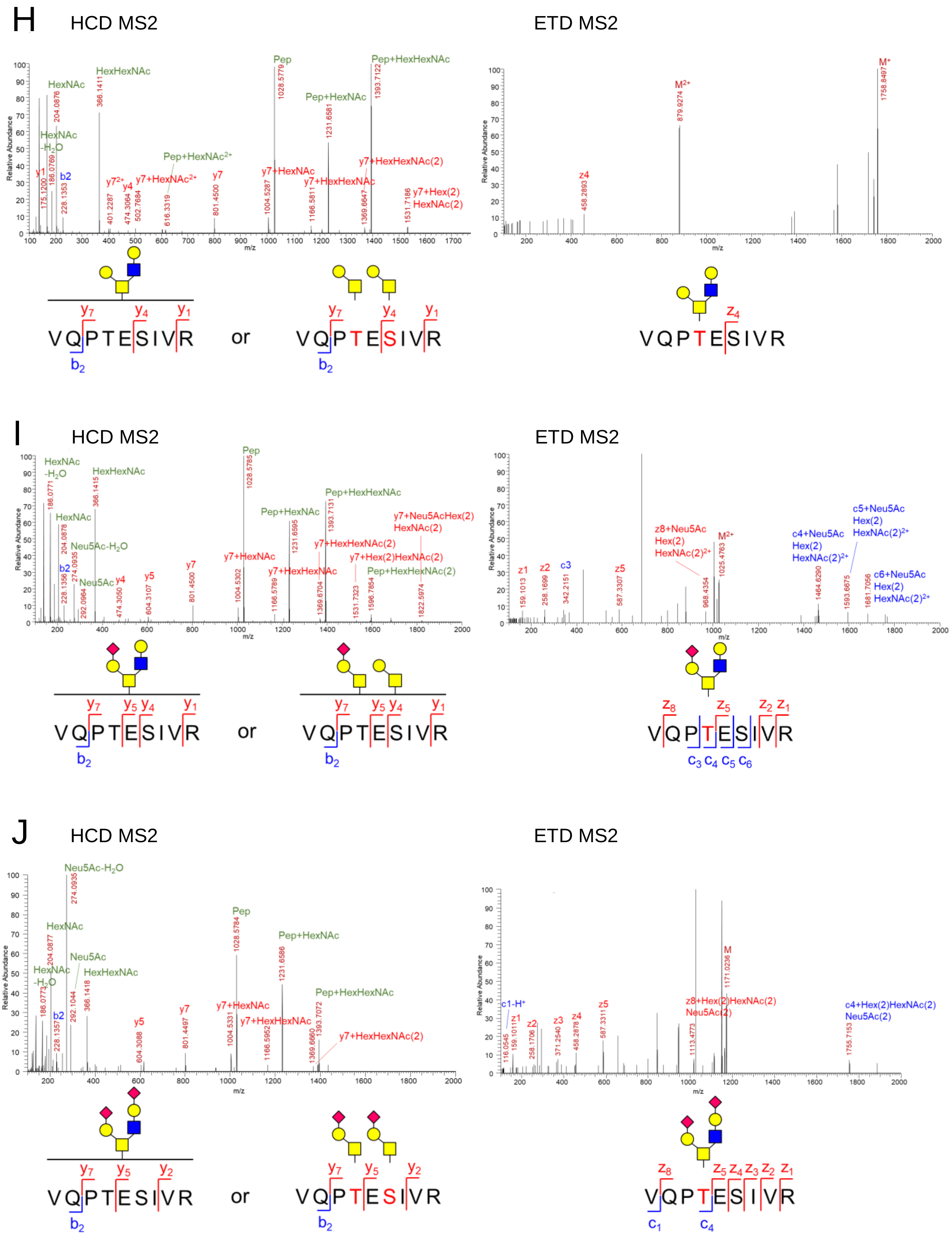

H
HCD MS2
ETD MS2
HexNAc
HexHexNAc
HexNAc -H2O
Pep
Pep+HexHexNAc
Pep+HexNAc
Pep+HexNAc2+
y7+HexNAc
y7+HexHexNAc
y7+HexHexNAc(2)
y7+Hex(2) HexNAc(2)
M2+
M+
z4
or
VQPTESIVR
I
HCD MS2
ETD MS2
Neu5Ac-H2O
Neu5Ac
y7+Neu5AcHex(2) HexNAc(2)
y7+Hex(2)HexNAc(2)
Pep+HexHexNAc(2)
z8+Neu5Ac Hex(2) HexNAc(2)2+
c4+Neu5Ac Hex(2) HexNAc(2)2+
c5+Neu5Ac Hex(2) HexNAc(2)2+
c6+Neu5Ac Hex(2) HexNAc(2)2+
J
HCD MS2
ETD MS2
z8+Hex(2)HexNAc(2) Neu5Ac(2)
c4+Hex(2)HexNAc(2) Neu5Ac(2)
c1-H+
M

**Figure S5 Representative HCD and ETD MS/MS spectra. (Related to Figure 1, 3 and S2)**
C. Representative HCD and ETD MS/MS spectra of intact O-glycopeptides of recombinant S protein with determined O-glycosites.

## S31

HCD MS2

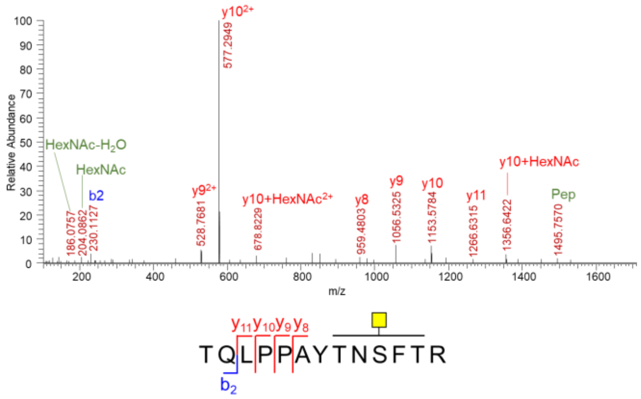


ETD MS2

Not detected

This site was confirmed in the MS/MS spectrum of S-ECD-3×FLAG protein in Fig. S6.

## T315 & S316

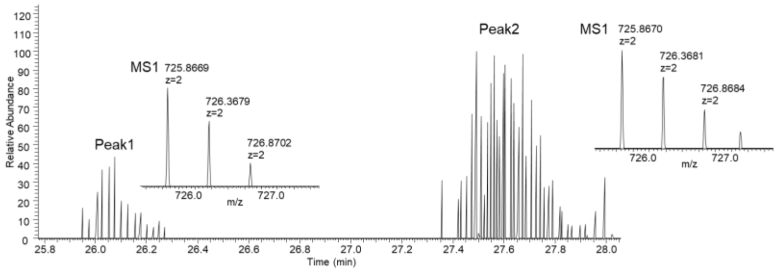


The extracted ion chromatogram and isotope distribution of O-glycopeptide $^{310}$GIYQTSNFR$^{319}$ with core 1 type GalNAcGal glycan.

Peak1

HCD MS2

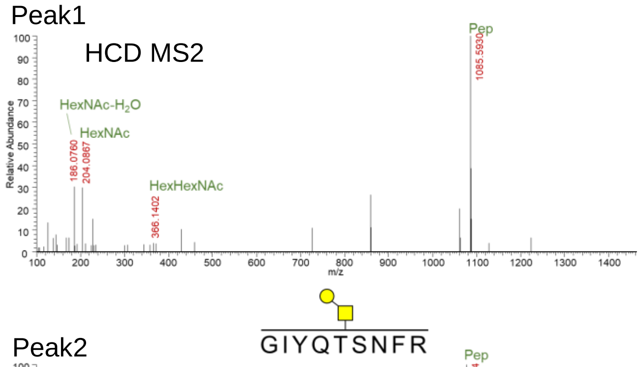


ETD MS2

Not detected

Peak2

HCD MS2

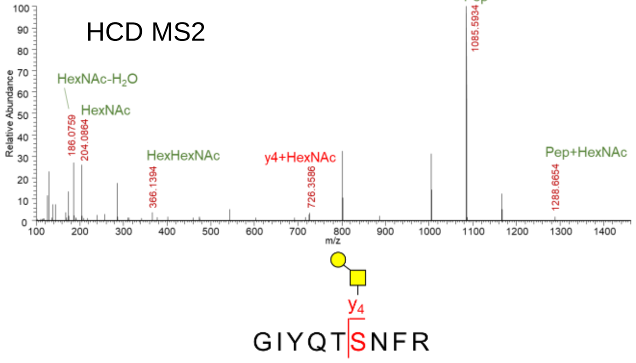


ETD MS2

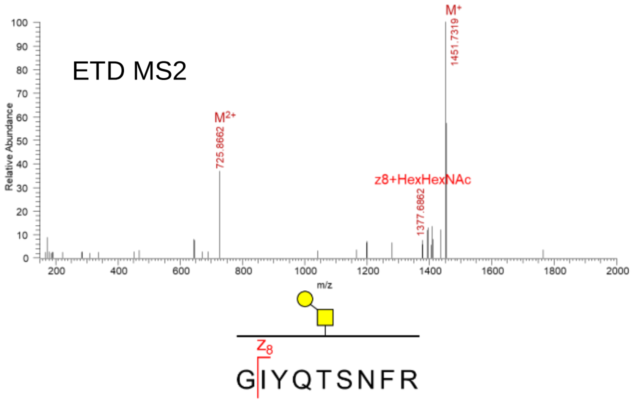


The glycosite of the O-glycopeptide in Peak2 is S316. Thus, the glycosite of the O-glycopeptide in Peak1 is T315.

# T323

HCD MS2

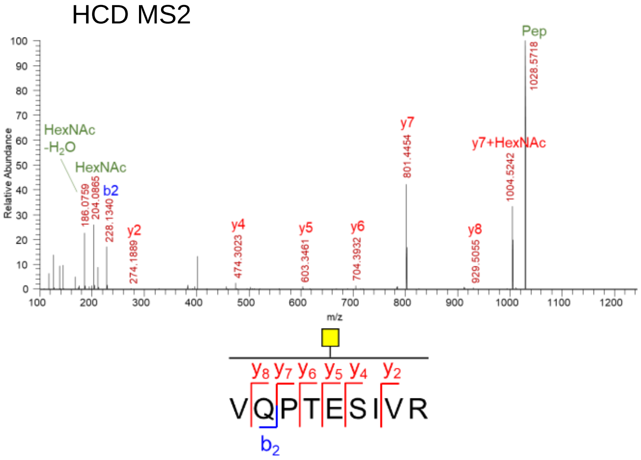


ETD MS2

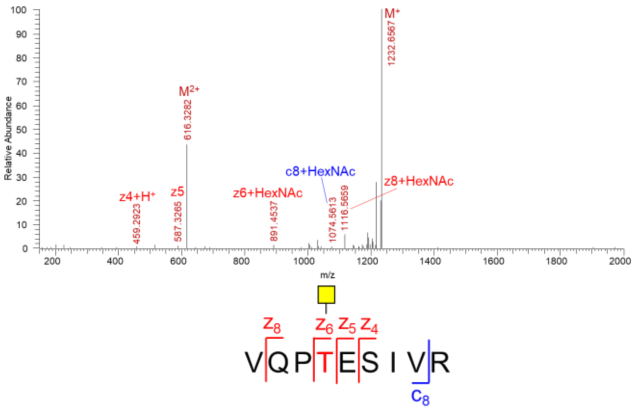


# T638

HCD MS2

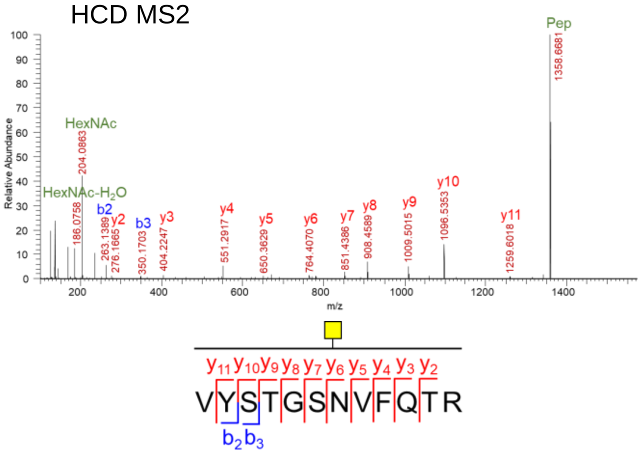


ETD MS2

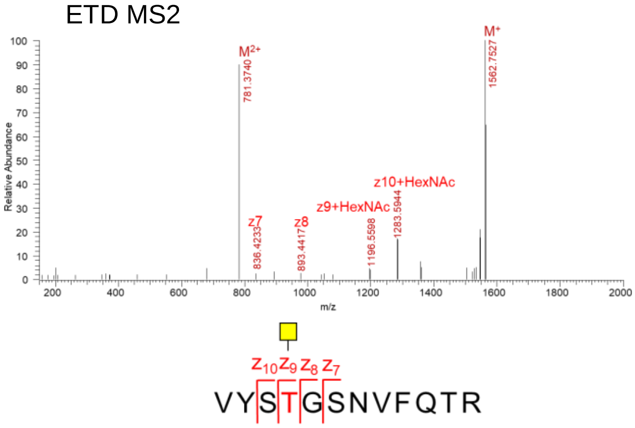


# S640

HCD MS2

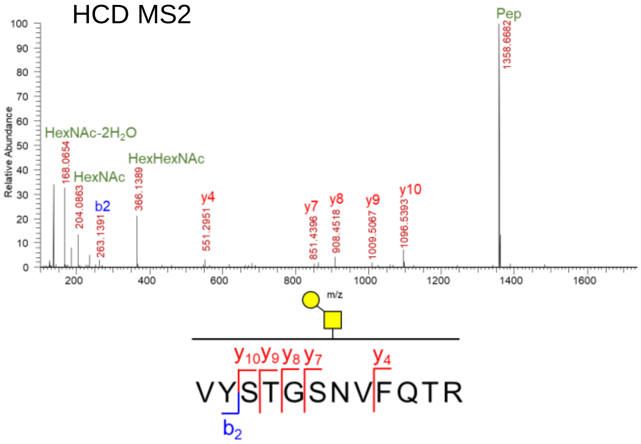


ETD MS2

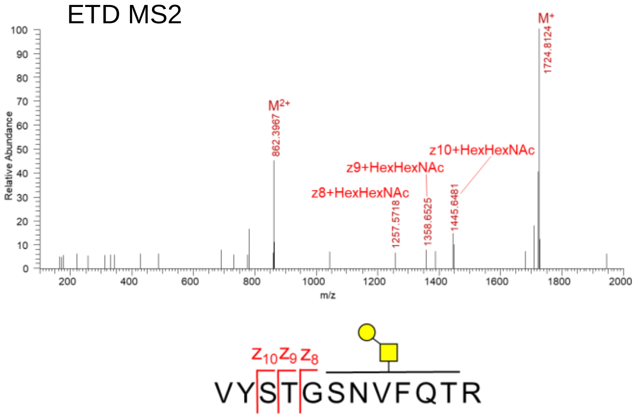


This site was confirmed in the MS/MS spectrum of S-ECD-3×FLAG protein in Fig. S6.

# T912

HCD MS2

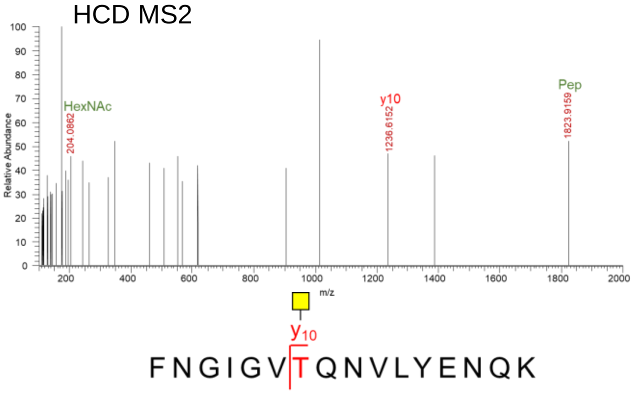


ETD MS2

Not detected

# S939 or S940 or T941 or S943

HCD MS2

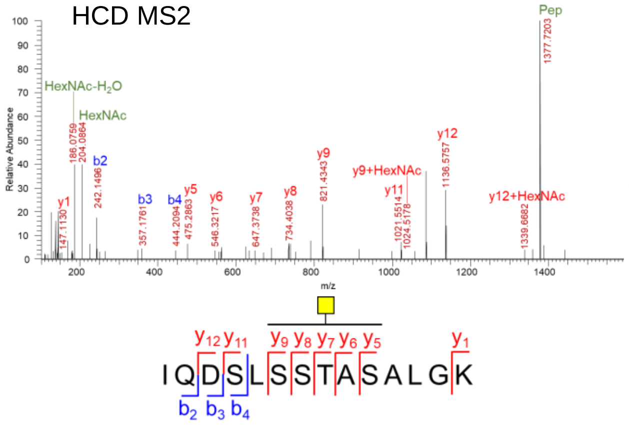


ETD MS2

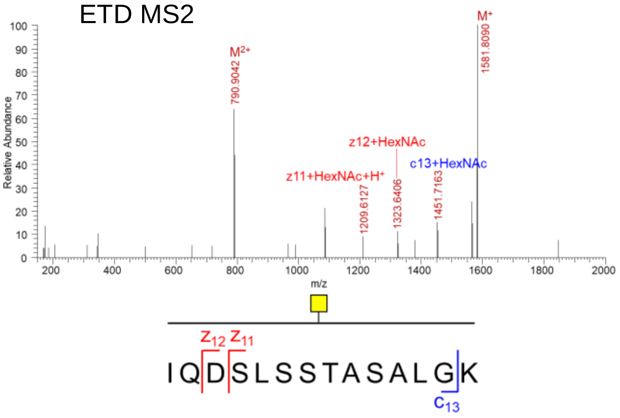


# T1027

HCD MS2

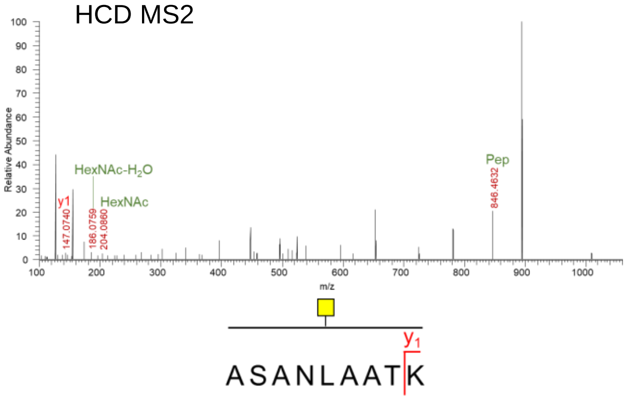


ETD MS2

Not detected

This site was confirmed in the MS/MS spectrum of S-ECD-3×FLAG protein in Fig. S6.

**Figure S5 Representative HCD and ETD MS/MS spectra. (Related to Figure 1, 3 and S2)**

D. Representative HCD and ETD MS/MS spectra of intact O-glycopeptides of recombinant S protein purified from ppGalNAc-T6 overexpressed HEK-293T cells.

# T29 & S31

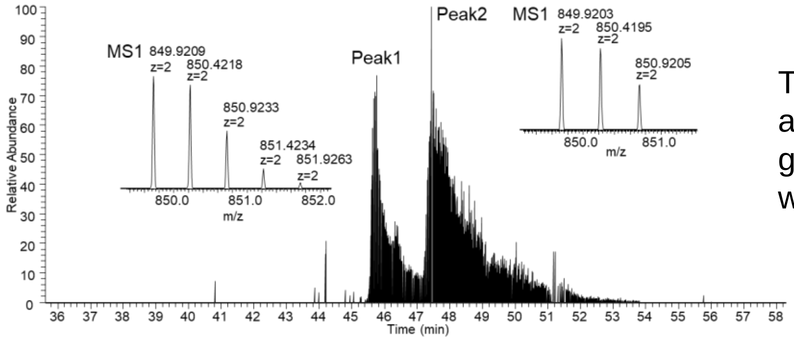


The extracted ion chromatogram and isotope distribution of O-glycopeptide [22]TQLPPAYTNSFTR[34] with Tn structure O-glycan.

Peak1 S31

HCD MS2

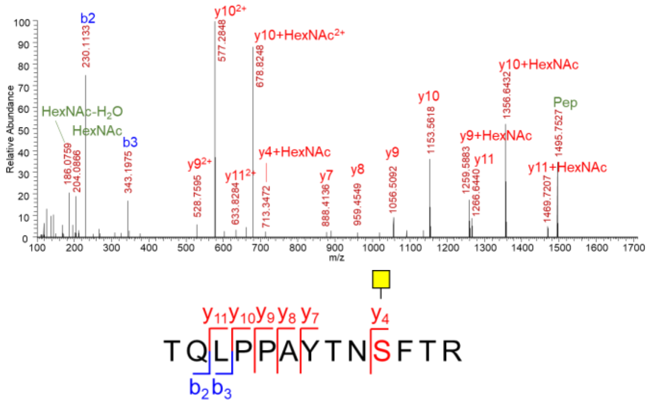


ETD MS2

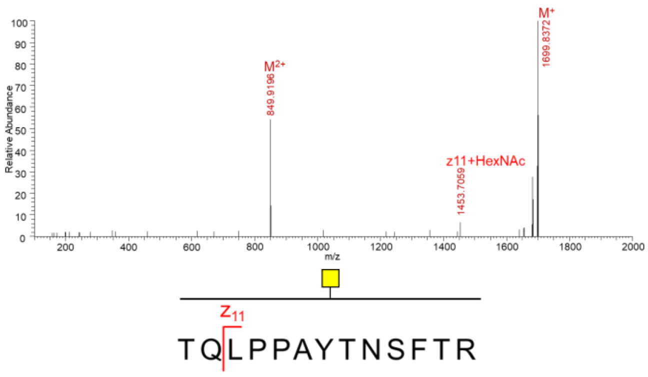


Peak2 T29

HCD MS2

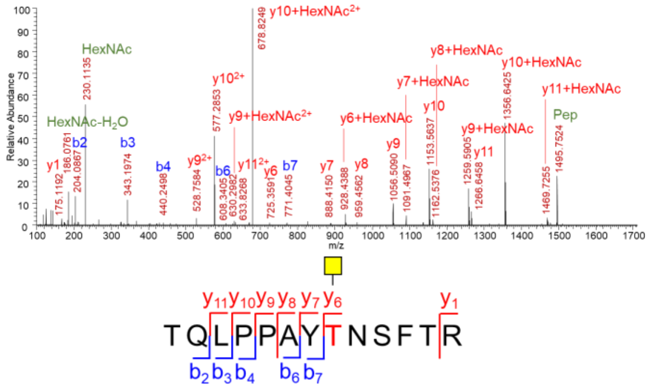


ETD MS2

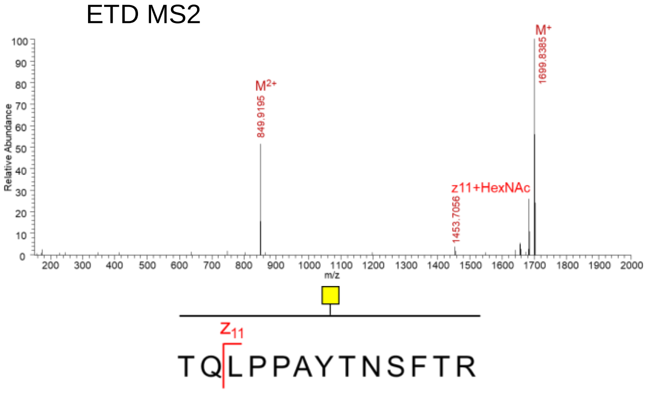


The glycosite of the O-glycopeptide in Peak1 is S31. Thus, the glycosite of the O-glycopeptide in Peak2 is T29.

# T307

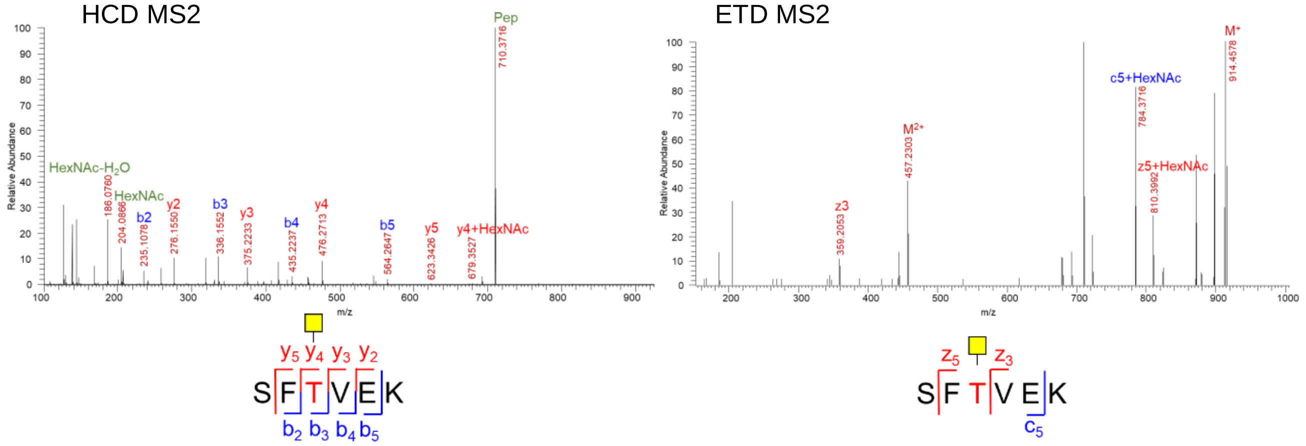


# T315 & S316

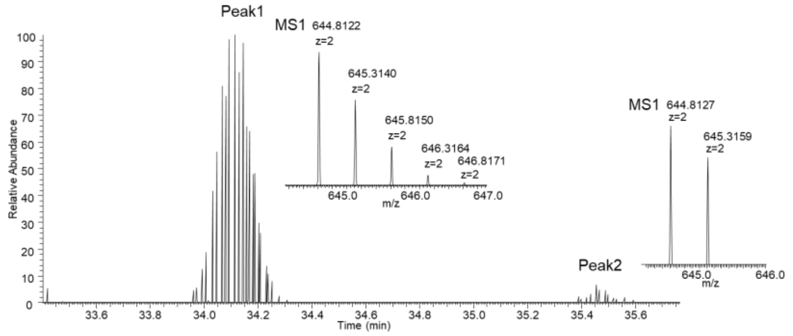


The extracted ion chromatogram and isotope distribution of O-glycopeptide $^{310}$GIYQTSNFR$^{319}$ with Tn structure O-glycan.

Peak1

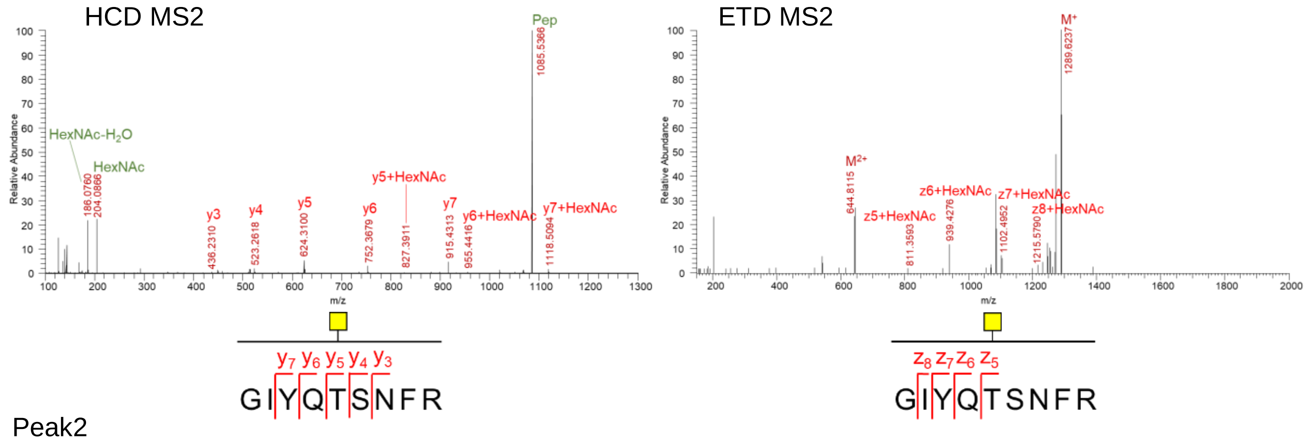


Peak2

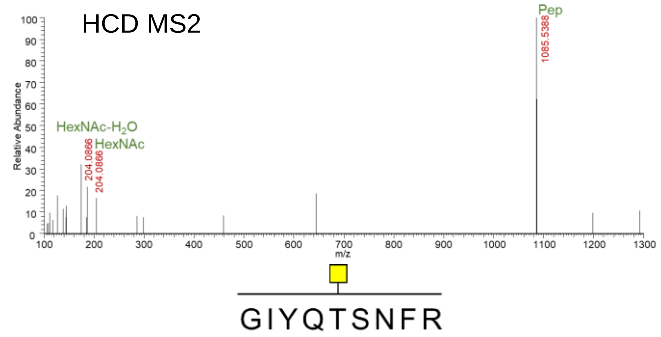


ETD MS2

Not detected

According to the results in Fig. S5, the glycosite of the O-glycopeptide in Peak1 is S315, and the glycosite of the O-glycopeptide in Peak2 is T316.

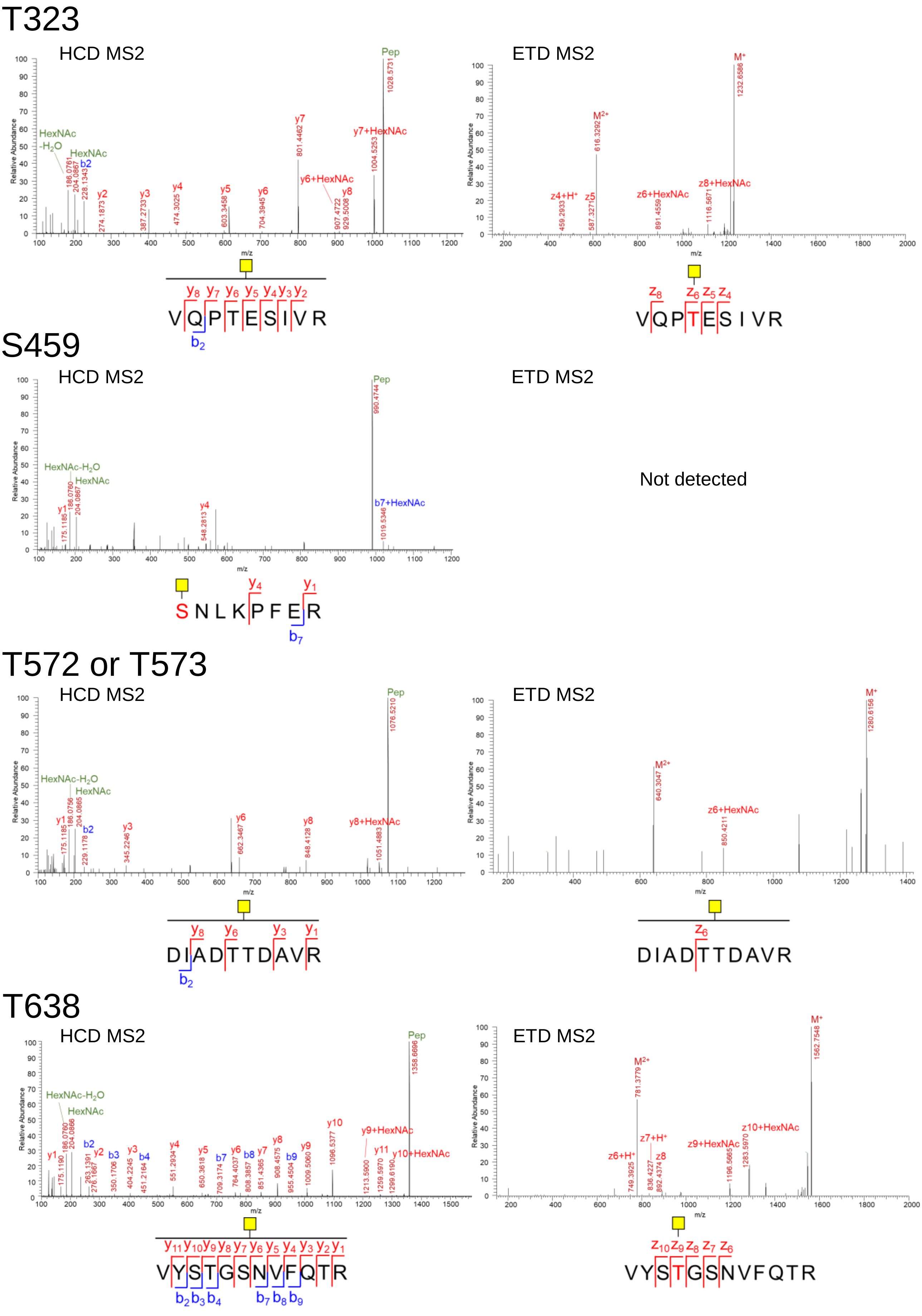

T323
HCD MS2
Pep
1028.5731
HexNAc -H2O
HexNAc
b2
186.0761
204.0867
228.1343
y2
274.1873
y3
387.2733
y4
474.3025
y5
603.3458
y6
704.3945
y7
801.4462
y6+HexNAc
y8
907.4722
929.5008
y7+HexNAc
1004.5253
ETD MS2
M+
1232.6586
M2+
616.3292
z4+H+
459.2933
z5
587.3271
z6+HexNAc
891.4559
z8+HexNAc
1116.5671
VQPTESIVR
S459
HCD MS2
Pep
990.4744
HexNAc-H2O
HexNAc
y1
175.1185
186.0760
204.0867
y4
548.2813
b7+HexNAc
1019.5346
ETD MS2
Not detected
SNLKPFER
T572 or T573
HCD MS2
Pep
1076.5210
HexNAc-H2O
HexNAc
y1
175.1185
186.0756
204.0865
b2
229.1178
y3
345.2246
y6
662.3467
y8
848.4128
y8+HexNAc
1051.4883
ETD MS2
M+
1280.6156
M2+
640.3047
z6+HexNAc
850.4211
DIADTTDAVR
T638
HCD MS2
Pep
1358.6696
HexNAc-H2O
HexNAc
y1
175.1190
186.0760
204.0866
b2
y2
263.1391
276.1667
b3
350.1706
y3
404.2245
b4
451.2164
y4
551.2934
y5
650.3618
b7
709.3174
y6
764.4037
b8
808.3857
y7
851.4365
y8
908.4575
b9
955.4504
y9
1009.5060
y10
1096.5377
y9+HexNAc
y11
1213.5900
1259.5970
y10+HexNAc
1299.6190
ETD MS2
M+
1562.7548
M2+
781.3779
z6+H+
749.3925
z7+H+
z8
836.4227
892.4374
z9+HexNAc
1196.5665
z10+HexNAc
1283.5970
VYSTGSNVFQTR
Relative Abundance
m/z

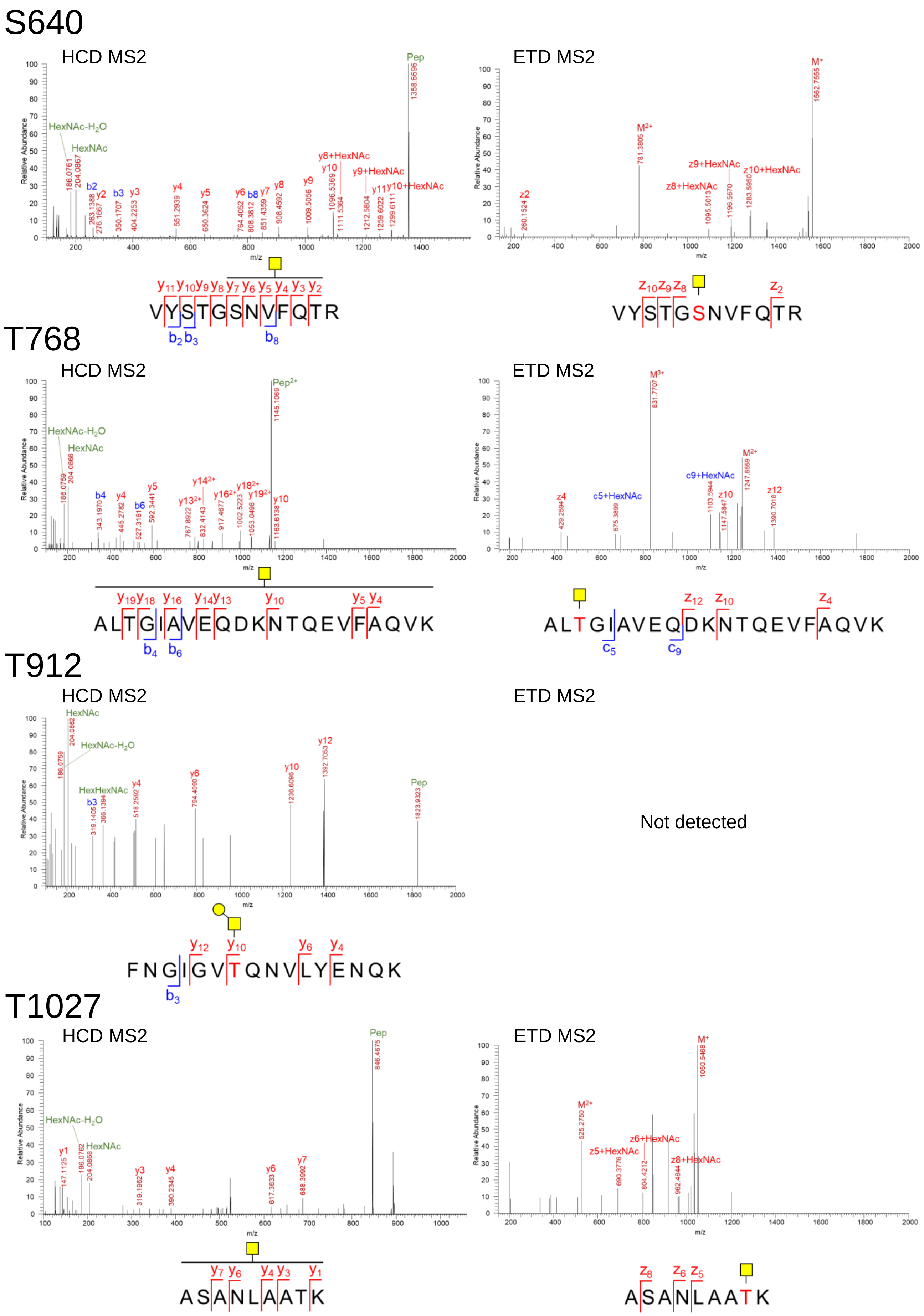

S640
HCD MS2
ETD MS2
VYSTGSNVFQTR
T768
HCD MS2
ETD MS2
ALTGIAVEQDKNTQEVFAQVK
T912
HCD MS2
ETD MS2
Not detected
FNGIGVTQNVLYENQK
T1027
HCD MS2
ETD MS2
ASANLAATK
Relative Abundance
m/z
HexNAc-H2O
HexNAc
HexHexNAc
Pep
Pep2+
M+
M2+
M3+

**Table S1. Detail LC-MS/MS information of all the O-glycopeptides detected in the three samples of S-ECD, S-ECD co-expressed with T6, and S1. (Related to Figure 1, 3 and S2)**

| Peptide | Glycan | Sites | Theoretical MW (Da) | Experimental MW (Da) | | | Retention time (min) | | |
|---|---|---|---|---|---|---|---|---|---|
| | | | | S | S&T6 | S1 | S | S&T6 | S1 |
| [R].VQPTESIVR.[F] | HexNAc | T323 | 1231.65 | 1231.65 | 1231.65 | 1231.66 | 28.90 | 29.45 | 29.15 |
| [R].VQPTESIVR.[F] | HexNAcHex | T323 | 1393.71 | 1393.70 | 1393.71 | 1393.71 | 27.14 | 27.35 | 26.99 |
| [R].VQPTESIVR.[F] | HexNAcHexNeuAc(2) | T323 | 1975.90 | — | — | 1975.91 | — | — | 37.24 |
| [R].VQPTESIVR.[F] | HexNAcHexNeuAc | T323 | 1684.80 | 1684.80 | 1684.80 | 1684.81 | 32.38 | 32.05 | 32.56 |
| [R].VQPTESIVR.[F] | HexNAc(2)Hex | T323 | 1596.79 | 1596.78 | 1596.79 | 1596.79 | 26.41 | 26.86 | 26.37 |
| [R].VQPTESIVR.[F] | HexNAc(2)Hex(2) | T323 | 1758.84 | 1758.84 | 1758.84 | 1758.85 | 26.43 | 26.76 | 26.28 |
| [R].VQPTESIVR.[F] | HexNAc(2)HexNeuAc | T323 | 1887.88 | — | — | 1887.89 | — | — | 31.62 |
| [R].VQPTESIVR.[F] | HexNAcNeuAc | T323 | 1522.75 | — | — | 1522.76 | — | — | 35.16 |
| [R].VQPTESIVR.[F] | HexNAc(2)HexNeuAc | T323 | 1887.88 | — | — | 1887.89 | — | — | 31.62 |
| [R].VQPTESIVR.[F] | HexNAc(2)Hex(2)NeuAc | T323 | 2049.93 | — | — | 2049.94 | — | — | 31.40 |
| [R].VQPTESIVR.[F] | HexNAc(2)Hex(2)NeuAc(2) | T323 | 2341.03 | — | — | 2341.07 | — | — | 35.69 |
| [R].VYSTGSNVFQTR.[A] | HexNAc | T638 | 1561.75 | 1561.76 | 1561.75 | 1561.75 | 37.79 | 38.26 | 39.05 |
| [R].VYSTGSNVFQTR.[A] | HexNAcHex | T638 | 1723.80 | 1723.80 | 1723.80 | 1723.81 | 37.47 | 37.72 | 37.62 |
| [R].VYSTGSNVFQTR.[A] | HexNAcHexNeuAc | T638 | 2014.90 | — | 2014.90 | 2014.86 | — | 42.64 | 41.46 |
| [R].VYSTGSNVFQTR.[A] | HexNAcHexNeuAc(2) | T638 | 2305.99 | — | — | 2306.02 | — | — | 29.56 |
| [R].VYSTGSNVFQTR.[A] | HexNAc(2)HexNeuAc | T638 | 2217.98 | — | — | 2218.00 | — | — | 39.28 |
| [R].VYSTGSNVFQTR.[A] | HexNAc(2)Hex(2)NeuAc | T638 | 2380.03 | — | — | 2380.00 | — | — | 36.76 |
| [R].VYSTGSNVFQTR.[A] | HexNAc(2)Hex(2)NeuAc(2) | T638 | 2671.13 | — | — | 2671.14 | — | — | 46.20 |
| [R].VYSTGSNVFQTR.[A] | HexNAc(2)Hex | T638 | 1926.88 | — | 1926.89 | — | — | 35.39 | — |
| [K].SFTVEK.[G] | HexNAc | T307 | 913.45 | — | 913.45 | 912.42 | — | 27.67 | 27.62 |
| [K].SFTVEK.[G] | HexNAcHex | T307 | 1075.50 | — | 1075.50 | 1075.55 | — | 26.16 | 26.16 |
| [K].SFTVEK.[G] | HexNAc(2)Hex | T307 | 1278.58 | — | — | 1278.59 | — | — | 26.00 |
| [K].SFTVEK.[G] | HexNAc(2)HexNeuAc | T307 | 1569.68 | — | — | 1569.69 | — | — | 31.89 |
| [K].SFTVEK.[G] | HexNAc(2)Hex(2)NeuAc | T307 | 1731.73 | — | — | 1731.74 | | — | 31.83 |
| [K].GIYQTSNFR.[V] | HexNAc | T315 | 1288.62 | 1288.67 | 1288.62 | 1288.62 | 27.60 | 34.11 | 34.30 |
| [K].GIYQTSNFR.[V] | HexNAcHex | T315 | 1450.67 | 1450.73 | 1450.73 | 1450.68 | 26.07 | 33.39 | 33.50 |
| [K].GIYQTSNFR.[V] | HexNAcHexNeuAc | T315 | 1741.77 | — | 1741.77 | 1741.77 | — | 39.46 | 37.36 |
| [K].GIYQTSNFR.[V] | HexNAcHexNeuAc(2) | T315 | 2032.86 | — | — | 2032.87 | — | — | 44.10 |
| [K].GIYQTSNFR.[V] | HexNAc(2)Hex | T315 | 1653.75 | — | — | 1653.76 | — | — | 30.99 |
| [K].GIYQTSNFR.[V] | HexNAc(2)HexNeuAc | T315 | 1944.84 | — | — | 1944.85 | — | — | 37.49 |
| [K].GIYQTSNFR.[V] | HexNAc(2)Hex(2) | T315 | 1815.80 | — | — | 1815.81 | — | — | 30.29 |
| [K].GIYQTSNFR.[V] | HexNAc(2)Hex(2)NeuAc | T315 | 2106.90 | — | — | 2106.91 | — | — | 36.62 |
| [K].GIYQTSNFR.[V] | HexNAc(2)Hex(2)NeuAc(2) | T315 | 2397.99 | — | — | 2398.01 | — | — | 41.54 |
| [K].GIYQTSNFR.[V] | HexNAc | S316 | 1288.62 | 1288.67 | 1288.62 | — | 29.93 | 35.46 | — |
| [K].GIYQTSNFR.[V] | HexNAcHex | S316 | 1450.67 | 1450.72 | 1450.73 | — | 28.02 | 26.30 | — |
| [K].GIYQTSNFR.[V] | HexNAcHexNeuAc | S316 | 1741.77 | — | 1741.77 | — | — | 35.42 | — |
| [R].VQPTESIVR.[F] | HexNAc | T325 | 1231.65 | — | — | 1231.66 | — | — | 29.39 |
| [R].VQPTESIVR.[F] | HexNAcNeuAc | T325 | 1522.75 | — | — | 1522.76 | — | — | 35.67 |
| [R].TQLPPAYTNSFTR.[G] | HexNAc | T29 | 1698.83 | — | 1698.83 | — | — | 47.43 | — |

Table S1. continued

| | | | | | | | | | |
|---|---|---|---|---|---|---|---|---|---|
| [R].TQLPPAYTNSFTR.[G] | HexNAc | S31 | 1698.83 | 1698.83 | 1698.83 | | 44.91 | 45.75 | — |
| [K].SNLKPFER.[D] | HexNAc | S459 | 1193.62 | — | 1192.55 | 1191.57 | — | 39.80 | 40.73 |
| [R].DIADTTDAVR.[D] | HexNAc | T572 or T573 | 1279.60 | — | 1279.60 | 1279.61 | — | 29.10 | 29.35 |
| [R].VYSTGSNVFQTR.[A] | HexNAc | S640 | 1561.75 | — | — | — | 38.01 | 38.42 | — |
| [R].VYSTGSNVFQTR.[A] | HexNAcHex | S640 | 1723.80 | — | — | — | 37.45 | 37.71 | — |
| [R].ALTGIAVEQDKNTQ EVFAQVK.[Q] | HexNAc | T768 | 2492.29 | — | 2493.30 | — | — | 54.84 | — |
| [R].FNGIGVTQNVLYEN QK.[L] | HexNAc | T912 | 2027.01 | 2027.01 | 2027.01 | — | 52.01 | 52.08 | — |
| [R].FNGIGVTQNVLYEN QK.[L] | HexNAcHex | T912 | 2189.06 | — | 2190.06 | — | — | 49.78 | — |
| [K].IQDSLSSTASALGK.[L] | HexNAc | S939 or S940 or T941 or S943 | 1580.80 | 1580.80 | 1580.80 | — | 36.33 | 36.55 | — |
| [R].ASANLAATK.[M] | HexNAc | T1027 | 1049.55 | 1049.52 | 1049.55 | — | 20.28 | 20.23 | — |
